# A bottom-up approach to understanding ultrasound mediated blood–brain barrier opening using long and short pulses

**DOI:** 10.1101/2025.06.11.659100

**Authors:** Yuanyuan Shen, Yicong Cai, Chaofeng Qiao, Zhihui Liu, Yiluo Xu, Mengni Hu, Ji Zhang, Lin Ma, Fenfang Li

## Abstract

Ultrasound-mediated blood-brain barrier (BBB) opening is promising for non-invasive, localized, and reversible drug delivery to the brain. However, the underlying mechanisms remain unclear, particularly at microscopic level in real time. Recently developed short-pulse ultrasound enhances safety by reducing side effects compared to conventional long-pulse ultrasound. Using vessel-mimicking microchannels cultured with brain endothelial monolayers, we directly observed bubble dynamics and cellular bioeffects under flow conditions. Intriguingly, cyclic jetting during stable cavitation occurs at long-pulse mode, accompanied by a few localized cell detachments and extensive sonoporation. In contrast, short-pulse ultrasound induced milder, uniform bubble dynamics, reversible sonoporation and calcium signaling, promoting safer BBB modulation. In vivo two-photon imaging and histology in mice confirmed these findings, showing that long-pulse ultrasound enabled higher drug delivery efficiency but caused localized endothelial damage, while short-pulse ultrasound facilitated uniform delivery and faster BBB recovery. The differential bubble dynamics and cellular responses correlated well between in vitro and in vivo models. This study establishes a cross-scale framework for real-time analysis of ultrasound-mediated BBB opening, revealing key biophysical factors governing safety and efficacy. The findings provide guidance for the optimization of ultrasound protocols for vascular drug delivery, with potential applications in treating brain tumors, neurodegenerative diseases, and other disorders.

## Introduction

The blood–brain barrier (BBB) is composed of tightly sealed brain vascular endothelial cells (ECs) that closely interact and communicate with other components such as astrocytes, pericytes, perivascular microglia and neurons, together strengthening the barrier functions and maintaining brain homeostasis [1, 2]. BBB tightly regulates the trafficking of molecules between the blood and the brain and protects the central nervous system (CNS) from toxins and pathogens [3]. However, the lower rate of transcytosis compared to non-CNS vasculature and the presence of tight junctions between brain ECs also severely impede most therapeutic drugs for the treatment of CNS diseases, including brain cancers and neurological disorders [4–6]. Focused ultrasound (FUS) is an emerging technique that can noninvasively deliver acoustic energy to the brain through the intact skull and actuate microbubbles (MBs) oscillation in the focal region to transiently, locally and reversibly open the BBB. Targeted delivery of therapeutics to the CNS has been achieved with this technique [7], including chemotherapeutics [8, 9], antibodies [10, 11], cytokines [12], neurotropic factors [13, 14], adeno-associated viruses [15, 16], stem cells [17], immune cells [18]. This technology has advanced clinical trials in human patients worldwide for the intervention of various brain pathological conditions (e.g., glioblastoma, Parkinson’s disease, Alzheimer’s disease and amyotrophic lateral sclerosis) [19–23].

Despite promising clinical results, real-time control of the ultrasound dose delivered through the skull and monitoring of the cavitation activity is necessary as the range of pressures for efficient and safe BBB opening is very narrow [24]. Long-pulse ultrasound sequences with 10–23.5 ms pulses emitted at a slow rate (≤L10 Hz) are typically used in preclinical and clinical studies for BBB disruption [25, 26], but has also been associated with some adverse bioeffects, such as red blood cell extravasation [27], transient edema [28], and inflammation [25, 29]. While long-pulse ultrasound has been extensively studied and optimized for BBB opening, one strategy using short-pulse sequences has been developed as a promising alternative that broadens the design space of parameters toward a wider safety window while maintaining effective delivery [30–35]. Short-pulse ultrasound sequences, such as the rapid short-pulse (RaSP) sequence, delivers significantly less acoustic energy to the brain compared to long-pulse sequences, tends to induce smaller openings, resulting in minimal tissue damage, facilitating a more spatially uniform delivery and rapid restoration of BBB integrity within 10 minutes [30, 33]. However, the biophysical mechanisms by which microbubbles enhance the permeability of BBB in both approaches remain unclear, which is crucial for further optimizing acoustic parameters to improve efficiency and clinical applicability while minimizing associated damage [36, 37].

Proposed physical mechanisms by which microbubbles enhance biological barrier permeability include acoustic streaming [38], inertial jetting [39–42], cyclic jetting [43], normal impact pressure [44] and viscous shear stress [45]. The behaviors of microbubbles are altered when they are situated within small vessels and in flow conditions, and acoustic radiation force can displace bubbles to the vessel wall and induce bubble clusters or bubble coalescence due to secondary Bjerkness force [37, 46, 47]. RaSP sequences have been reported to produce spatiotemporally uniform cavitation distributions during flow conditions, and sustain microbubble activity for longer durations while fast microbubble destruction occurred for long pulses [48]. However, another study demonstrated persisting cavitation activity during long tone bursts [49]. Both studies employed passive cavitation detection and direct high-speed optical observation with limited spatiotemporal resolution. It is well established that stable or inertial cavitation of microbubbles could deliver drugs across cell monolayer through several pathways, including increased endocytosis [50], cell junction opening [51, 52], sonoporation [45] and direct cell detachment [39]. Besides, fluid flow and calcium signaling have also been found to play important roles for endothelium membrane permeabilization [41, 51, 53–55]. There is a lack of consensus that underscores the formidable challenge of directly observing bubble behavior and cellular bioeffects and correlating it with endothelial layer permeability change as well as *in vivo* BBB opening characteristics.

Therefore, in this study, we investigate the biophysical mechanisms underlying ultrasound-mediated BBB opening at long pulse and short pulse sequences from a bottom-up approach. We utilize *in vitro* vessel-mimicking microchannels to resolve bubble dynamics and cellular bioeffects at flow conditions and correlate the results with real-time *in vivo* 2-photon imaging for the dynamics and characteristics of BBB opening. This bottom-up approach enables us to uncover key insights into the biophysics of microbubble-mediated BBB permeabilization, which could guide the future developments of safer and more efficient BBB opening with ultrasound technology.

## Results

### Experimental Setup and Study Design

Microbubbles with a lipid shell and a core of perfluoropropane gas were synthesized following previously established protocols [56]. The freshly synthesized microbubbles displayed a polydisperse nature, with an average diameter of 1.4 μm and a concentration of 2.1 × 10^9^/mL (Fig. S1a). Ninety percent of the microbubbles had diameters below 2.0 μm (d90). Over five days following preparation, the microbubbles’ characteristics were monitored. By the fifth day, the mean diameter had increased by 15.8%, while the concentration decreased by 28.5%, indicating a relatively stable profile. A 1.125 MHz ring ultrasound transducer was carefully aligned with the vessel-mimicking microchannels (Fig. 1a) for studying both bubble dynamics and cellular bioeffects on an inverted optical microscope (Fig. S1b). The acoustic pressure output by the ring transducer was measured to be at its geometric center in the X-Y plane with a -6 dB diameter of around 1.5 mm after PDMS attenuation (Fig. 1b). Bubble dynamics were recorded with a 63X objective by high-speed imaging synchronized with the ultrasound pulses (Fig. S1 c-e). Cellular bioeffects were monitored by concurrent fluorescence imaging with propidium iodide (membrane poration indicator) and calcein (membrane integrity indicator) or Fluo-4 (Ca^2+^ signaling indicator) loaded in the cells (Fig. S1f). Long pulse and short pulse ultrasound sequences used in this study were shown in Fig.1c. The same ring transducer was used for two-photon intravital imaging capable of real-time observation of the BBB opening (Fig. 1d) with cranial window, which could observe brain vasculature at different depths from 0 μm to 600 μm (Fig. S2). Fig. 1e illustrates the two-dimensional mapping of the acoustic field emitted by the ring transducer, measured using a hydrophone at 0.5 mm below the cranial window coverslip.

**Figure 1.**
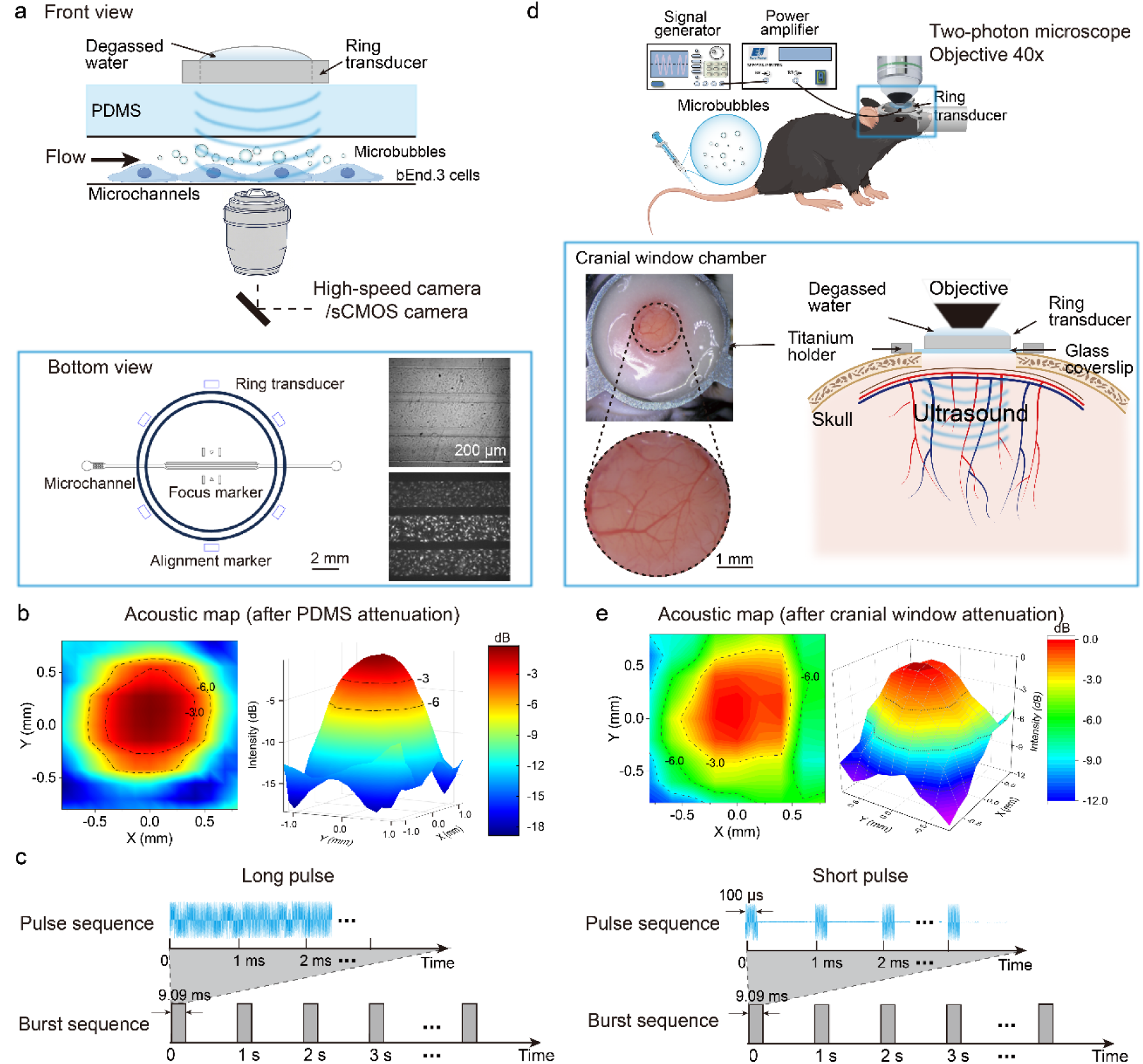
Experimental setup and design. (a) Schematic of the experimental setup for ultrasound stimulation, imaging of bubble dynamics and cellular bioeffects inside microchannels on an inverted microscope through a high-speed camera and an sCMOS camera, respectively. The enlarged bottom view shows the alignment of the ring ultrasound transducer with the microchannels, and the bright field and fluorescence images of the cell monolayer culture inside the microchannels. (b) Characterization of the acoustic field produced by the 1.125 MHz ring ultrasound transducer in the X-Y plane after PDMS attenuation. The acoustic pressure is shown in dB relative to the peak value. (c) Schematic of the ultrasound waveforms used in the experiments: long and short pulse modes. (d) Illustration of the two-photon intravital imaging setup capable of real-time observation of the BBB opening via ultrasound with microbubbles. The bottom inset shows a photograph of the cranial window and experimental apparatus. (e) Mapping of the acoustic field calibrated in water at a depth of 0.5 mm beneath the cranial window glass coverslip.

### Bubble Dynamics in Microchannels at Short Pulse and Long Pulse Ultrasound Sequences

The captured bubble dynamics inside the microchannel under short pulse and long pulse ultrasound exposure at different acoustic pressures are shown in Fig. 2. At short pulse of 0.25 MPa, where ultrasound lasted for 100 μs per pulse (0-100 µs), we observed displacement of microbubbles by acoustic radiation force and mild bubble coalescence due to secondary Bjerkness force (Fig. 2a, and Movie S1). At a higher acoustic pressure of 0.5 MPa, displacement and coalescence of microbubbles became more evident, resulting in larger bubble size and reduced number of bubbles (Fig. 2a, Movie S2). In comparison, at the long pulse mode of 0.25 MPa, where each ultrasound pulse lasted for 9.09 ms, displacement and clustering of microbubbles were observed (Fig. 2b, Movie S3). At higher acoustic pressure of 0.5 MPa in long pulse mode, after bubble displacement, clustering and coalescence in the early stage (0-1000 μs), both stable and inertial cavitation effects occurred in the later stage with bubble oscillation and collapse (1400-9160 μs) (Fig. 2b, Movie S4). Interestingly, some microbubbles underwent coalescence (S1) and then sustained oscillation (S2 and S3) at long pulse mode with the equilibrium bubble size resonant with the ultrasound center frequency (bottom two panels of Fig. 2b). Moreover, the formation of a jet propagating towards the substrate during the collapsing phase of the bubble was observed from 2500 to 3500 μs as indicated by the dark spot at the bubble center (bottom panel S3) and the temporal evolution of the volume-weighted bubble diameter (inset) of Fig. 2b (on the right). This cyclic jet during microbubble oscillation was reported by our previous study [57] (Fig. S10) and a recent report [43].

**Figure 2.**
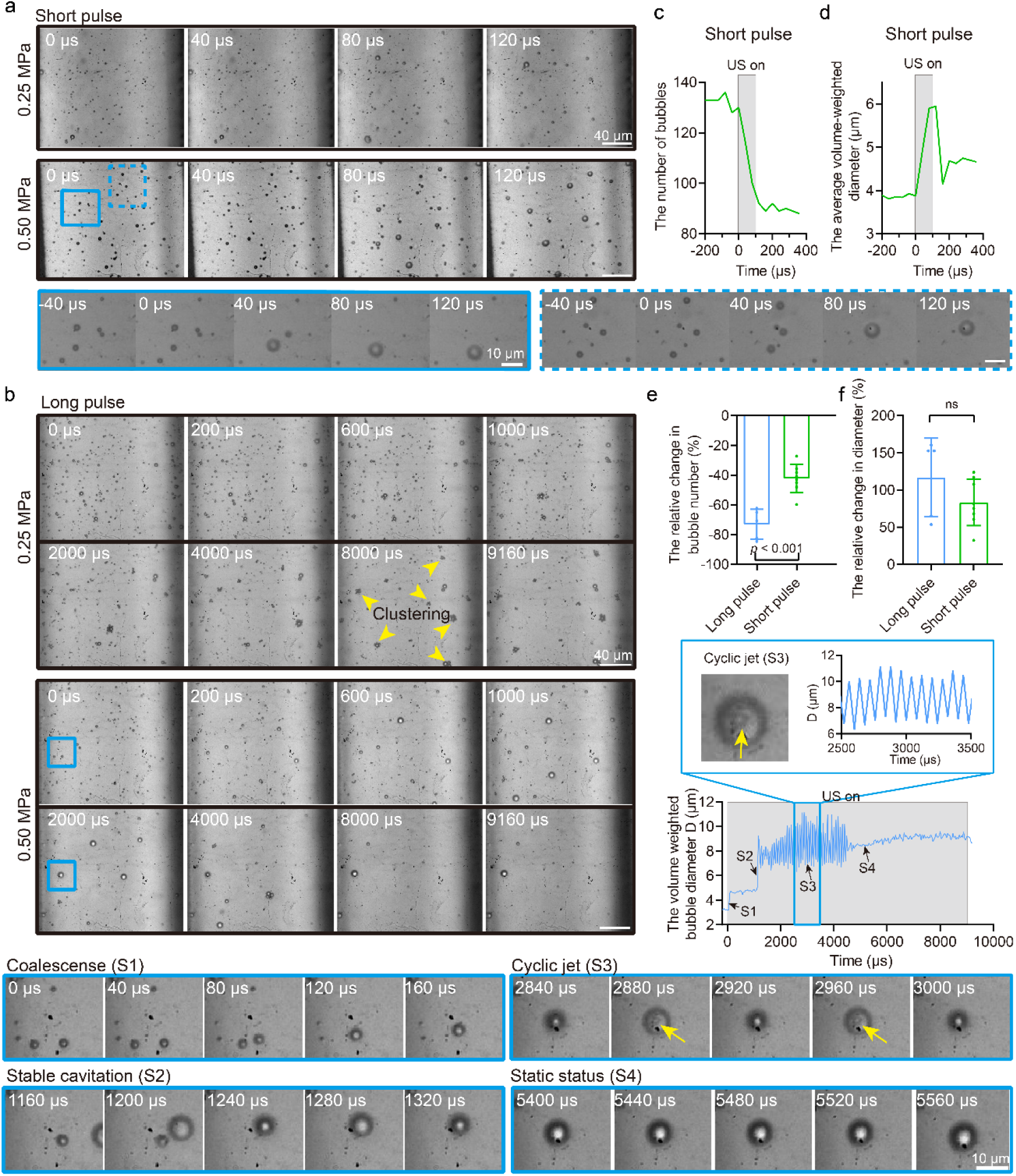
Bubble dynamics inside the microchannel under short pulse and long pulse ultrasound exposure at a flow rate of 75 μL/min under 0.25 and 0.5 MPa acoustic pressures. (a) High-speed recordings of the microbubbles inside the microchannel under short pulse ultrasound exposure (t=0-100 μs) at 0.25 and 0.5 MPa acoustic pressure. Bottom: Enlarged view of the displacement and coalescence of microbubbles inside the blue boxes in (a). (b) High-speed recordings of the microbubbles inside the microchannel under long pulse ultrasound exposure (t=0-9090 μs) at 0.25 and 0.5 MPa acoustic pressure. Bottom: Enlarged view of the oscillation of individual microbubbles inside the blue box in (b). The bubble kept oscillating from 1240 μs when generated from the coalescence of two bubbles till 4400 μs before it merged with another bubble (Movie S5). (c) Representative time evolution of the average volume weighted diameter of the bubbles in each image frame at short pulse mode under 0.5 MPa acoustic pressure. (d) Representative time evolution of the total number of bubbles in each image frame at short pulse mode under 0.5 MPa acoustic pressure. (e) and (f) Statistical analysis and comparisons of the bubble number reduction and relative bubble size increase between short pulse and long pulse mode at 0.5 MPa acoustic pressure. Each data point in panel e and f depicts the measurement from an independent microfluidic chip experiment. N=5 to 8 in panel e and f. The student t-test was used for statistical analysis.

Next, we calculated the total number of bubbles in each image frame and its variation with time (Fig. 2c). Similarly, we calculated the average volume-weighted diameter of the bubbles in each image frame and its time evolution (Fig. 2d). Based on these, the relative diameter change of the bubbles and the bubble number change can be obtained, see *In vitro* Image Processing and Data Analysis in the Materials and Methods section. Figure 2c shows that in short pulse mode, the total number of bubbles decreased after ultrasound exposure. The reduction was significantly higher at 0.5 MPa acoustic pressure (41%) than that (25%) at 0.25 MPa (Fig. S3a, left panel). The average volume-weighted diameter increased during ultrasound exposure (Fig. 2d), and the relative bubble diameter increase was more significant at a higher acoustic pressure of 0.5 MPa than 0.25 MPa (80% vs. 24%) (Fig. S3a, right panel). In long pulse mode, there was no significant difference in the bubble number changes between 0.25 and 0.5 MPa as both showed considerable reduction (60% and 75%) (Fig. S3b). The relative bubble diameter increase was more significant at higher acoustic pressure of 0.5 MPa in long pulse mode (115% vs. 45%). We further compared the difference between short and long pulse modes under the same acoustic pressure. A significant reduction in bubble numbers was found in long pulse mode compared to that in short pulse mode at both 0.25 MPa (Figures S3c (left panel) and 0.5 MPa (Fig. 2e). However, the relative bubble diameter increase was only significantly higher in long pulse mode at a lower acoustic pressure of 0.25 MPa (Fig. S3c, right panel) compared to that in short pulse mode, which might be due to acoustic pressure dependent bubble coalescence and the similar maximum expansion of the bubbles under 0.5 MPa (Fig. 2f).

### Cellular Bioeffects induced by Long Pulse and Short Pulse Ultrasound Exposure

Next, we investigated how the differential bubble dynamics would affect the cellular bioeffects in microchannels under long pulse and short pulse ultrasound exposure at 0.5 MPa at a flow rate of 75 uL/min, relevant to *in vivo* BBB opening. Cell detachment and compromise of monolayer morphology were observed after long pulse ultrasound treatment (red dashed box in Fig. S4a). In contrast, no discernable cell detachment was found after short pulse treatment (Fig. S4b).

To further dissect the dynamics of bubble-induced cellular bioeffects, including cell detachment, and membrane poration, we conducted concurrent fluorescence imaging of cellular PI uptake and calcein leakage to track cell membrane integrity. A representative recording in long pulse mode is shown in Figure 3a. Before ultrasound (-7 s), all the viable cells were loaded with calcein, and few cells showed PI uptake. When long pulse ultrasound was turned on, cellular calcein in several locations gradually decreased (Fig. 3b, left column, t=18-40 s and movie S6) while PI uptake occurred in some of these regions later (Fig. 3b, right column, t=63-192 s, and movie S6). After ultrasound exposure, the number of PI-positive cells gradually increased at the positions where the calcein became dark during (t=63-380 s). Cell detachment occurred at the region in the dashed box in Fig. 3a. Moreover, the detachment was a dynamic process, as could be seen from the white dashed lines in Fig. 3b, which tracked the upper right border of the detachment and moved gradually towards the upper right direction to enlarge the detachment (63-170 s). Cells nearby were also transferred in similar directions (the cell circled in Fig. 3b). For more details about the dynamics of cell detachment and membrane poration, please refer to supplementary movie S6. A representative fluorescence recording at short pulse mode is shown in Fig. S5 and Movie S7. During ultrasound exposure, cellular calcein also gradually decreased in a few locations but to a smaller extent. PI uptake gradually increased later at the positions where cellular calcein became dark. It is interesting to note that no cell detachment or discernable cell movement was found in short pulse mode, and the spatial distribution of cells showing PI uptake seems more uniform than long pulse mode. For more details about the dynamics of cell membrane poration at short pulse mode, please refer to supplementary movie S7.

**Figure 3.**
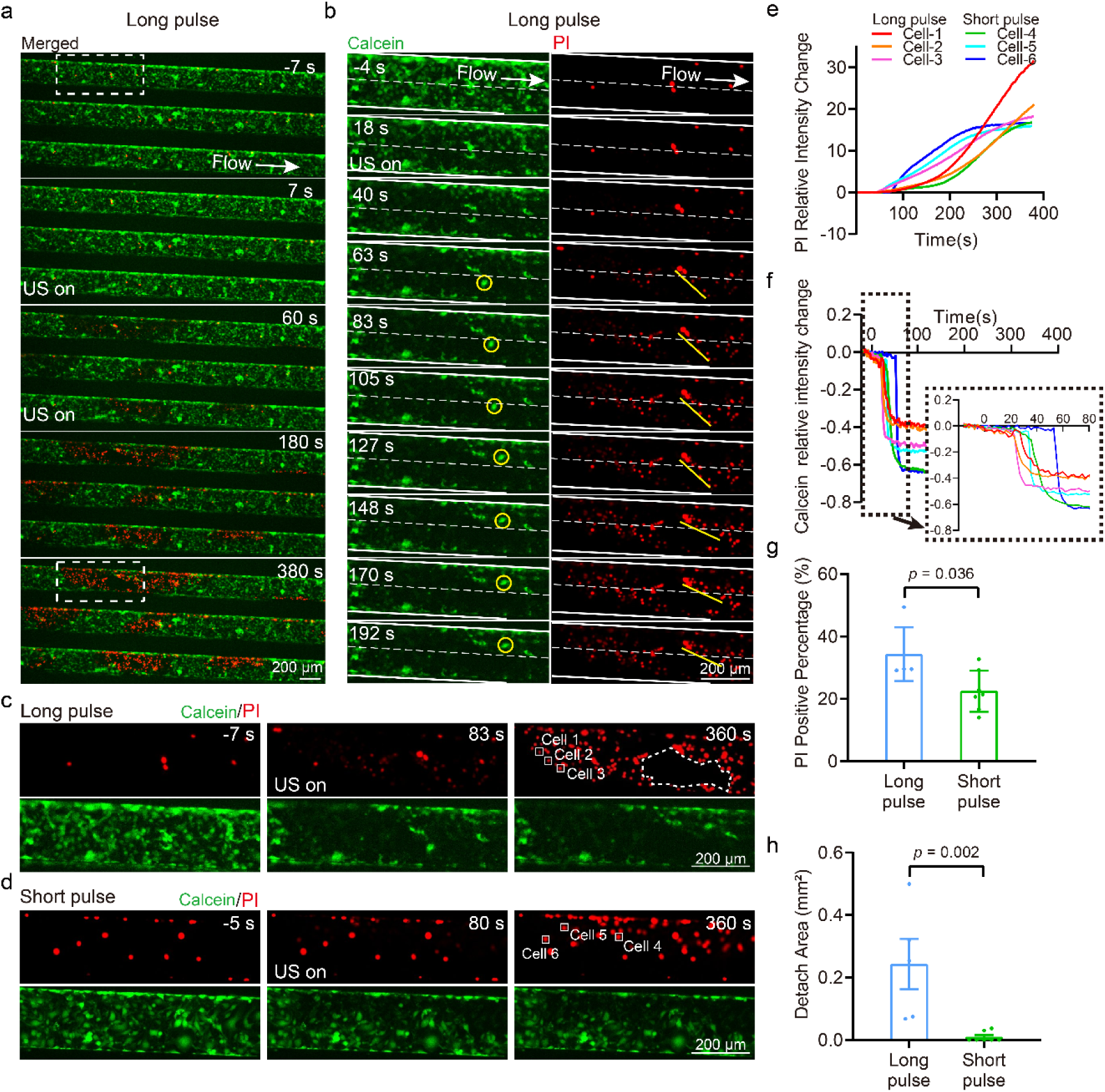
Characteristics of cell membrane poration and integrity change under 0.5 MPa ultrasound exposure. (a) Merged image sequence showing the calcein leakage (green) and PI uptake (red) in bEnd.3 cell monolayer in the microchannels before, during (0-60 s), and after 0.5 MPa ultrasound exposure at long pulse mode. (b) An enlarged view of the dashed box in (a) showing the time evolution of intracellular calcein (left), PI uptake (right) and cell detachment. The two split channels share the same time labels and scale bar. All the scale bars denote 200 μm. (c) and (d) PI uptake (red, upper panel) and calcein leakage (green, bottom panel) in bEnd.3 cell monolayer in the microchannels before (-7 s or -5 s), soon after (83 s or 80 s), and 360 s after 0.5 MPa ultrasound exposure at long and short pulse mode, respectively. The dashed line in panel c depicts the outline of the cell detachment region. The scale bar denotes 200 μm. (e) The relative PI uptake (I-I_0_)/I_0_)_PI_ vs. time for exemplary cells labeled by 1-3 at long pulse mode in (c) and 4-6 at short pulse mode in (d). (f) The relative calcein leakage (I-I_0_)/I_0_)_calcein_ vs. time for exemplary cells labeled by 1-3 at long pulse mode in (c) and 4-6 at short pulse mode in (d). The same color coding is used in (e) and (f). (g) and (h) The percentage of cells showing PI uptake and the total detachment area in each independent microfluidic chip experiment, respectively. N=5 and 7 for long pulse and short pulse mode, respectively. The student t-test was used for statistical analysis.

We took a closer look at the membrane integrity of the non-detached cells by their PI uptake and calcein leakage at both pulse sequences (Fig. 3c and 3d). Exemplary cells labeled by #1-3 at long pulse mode and #4-6 at short pulse mode demonstrated PI uptake after 60 s ultrasound exposure. Intriguingly, the relative PI intensity changes in cells #1-3 (long pulse mode) kept rising throughout the fluorescence recording, denoting irreversible membrane poration [58]. In contrast, the PI uptake in cells #5-6 and cell #4 (short pulse mode) reached a plateau around 250 s and 350 s, respectively, indicating resealing of the membrane pores (Fig. 3e). The averaged cellular calcein leakage in cell #1-3 (long pulse mode) was faster than that in cell # 4-6 (short pulse mode), likely due to a quicker comprise of the cell membrane by microbubble activities in long pulse mode (Fig. 3f). Nevertheless, they all reached plateau before t=75 s, which was faster than PI uptake due to their smaller stokes radii (0.7 nm vs. ∼1 nm). We further summarized and compared the percentage of cells showing PI uptake in each independent experiment at both modes. The results showed that PI positive cell percentage at long pulse mode was significantly higher than that at short pulse mode (Fig. 3g). Besides, the total detached area in each independent experiment was also considerably larger at long pulse mode while no evident cell detachment was observed at short pulse mode. These findings are consistent with our observations of the aforementioned bubble dynamics.

Recently, Ca^2+^ signaling has also been found to influence cell fate after sonoporation and regulate tight junction opening [41, 51, 55]. Therefore, we performed concurrent Ca^2+^ imaging and PI imaging for bEnd.3 monolayer culture in the vessel-mimicking channel. When exposed to long pulse ultrasound, notable Ca^2+^ response occurred at the sites of sonoporation, and Ca^2+^ signaling reached its peak then decayed before PI uptake was observed (Fig. S6). We also found cell detachment at some locations, where strong Ca^2+^ signaling was observed before PI uptake and cell detachment took place (Fig. S7). At short pulse mode, Ca^2+^ signaling was recorded at both mildly and highly sonoporated regions (Fig. 4 and Fig. S8). Overall, we found a more uniform Ca^2+^ signaling pattern inside the three microchannels at short pulse mode (Fig. 4a) compared to that of long pulse mode where Ca^2+^ signaling was more obvious at the locations of sonoporation or cell detachment (Fig. S6 and Fig. S7).

**Figure 4.**
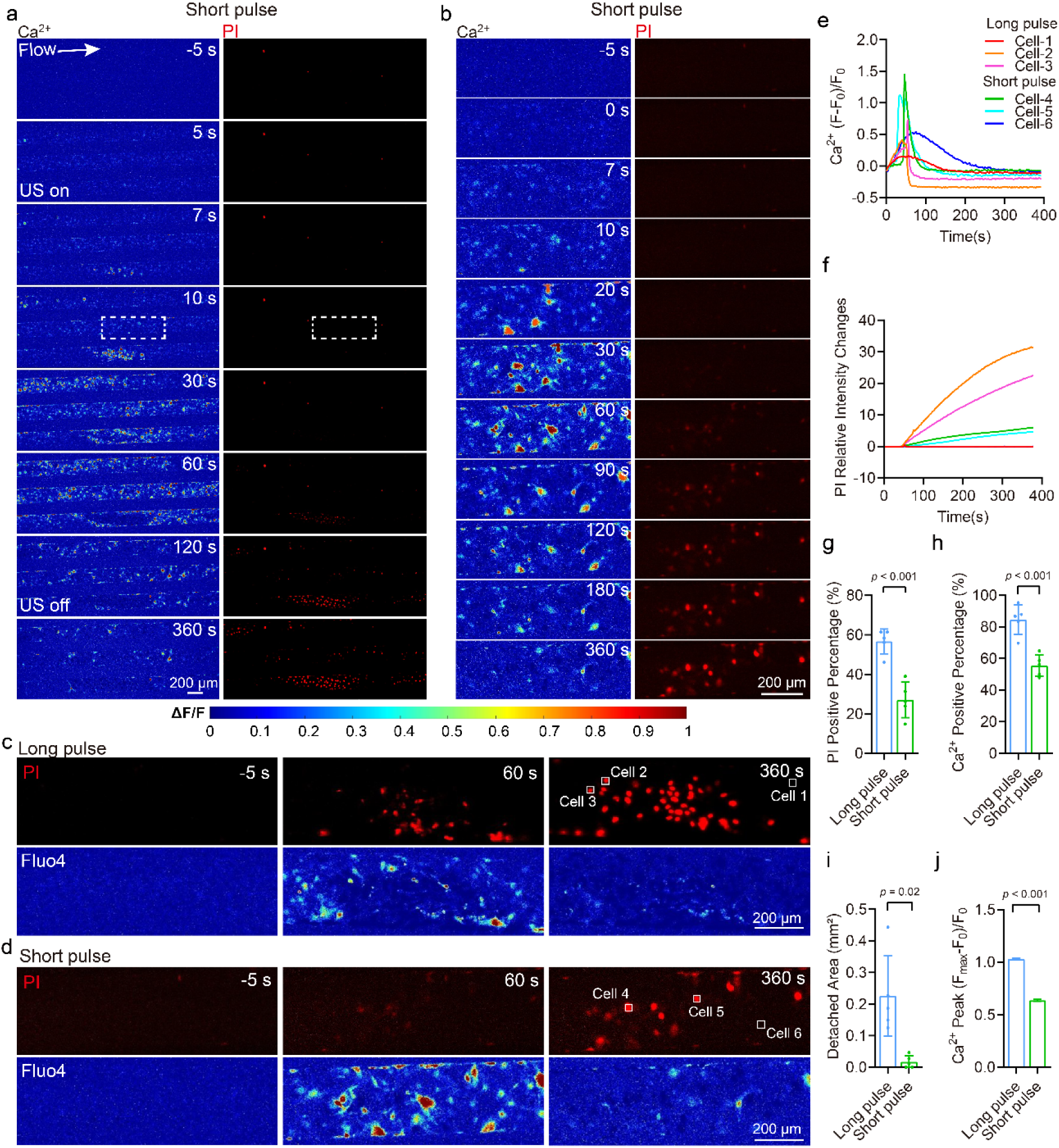
Characteristics of cellular Ca^2+^ signaling and membrane poration of bEnd.3 monolayer culture in the vessel-mimicking channel under 0.5 MPa ultrasound exposure. (a) Image sequences showing Ca^2+^ signaling (Fluo-4, pseudo color) and PI uptake (red) before, during (0-60 s), and after 0.5 MPa ultrasound exposure at short pulse mode. (b) Enlarged view of the dashed box in (a) showing the time evolution of Ca^2+^ signaling (left) and PI uptake (right). The two split channels share the same time labels and scale bar. (c) and (d) PI uptake (red, upper panel) and Ca^2+^ signaling (pseudo color, bottom panel) in bEnd.3 cell monolayer in the microchannels before (-5 s), during (60 s), and after (360 s) 0.5 MPa ultrasound exposure at long pulse and short pulse mode, respectively. The same pseudo color coding is used here as in panel a-b. (e) Ca^2+^ response ΔF/F vs. time for exemplary cells labeled by 1-3 at long pulse mode (c) and 4-6 at short pulse mode in (d). The relative PI uptake (I-I_0_)/I_0_)_PI_ vs. time for exemplary cells labeled by 1-3 at long pulse mode and 4-6 at short pulse mode is shown in (f). The same color coding for cells is used in panel e-f. (g) and (h) The percentage of cells showing PI uptake and Ca^2+^ signaling in each independent microfluidic chip experiment, respectively. (i) The total detachment area in each independent microfluidic chip experiment. Five independent microfluidic chip experiments were performed each for long and short pulse modes. (j) Ca^2+^ peak values were measured from individual cell nuclei in the above experiments. N=6275 and 5492 cells for long pulse and short pulse mode, respectively. The student t-test was used for statistical analysis in panels g-j.

We further examined the non-detached cells by their PI uptake and Ca^2+^ signaling at both pulse sequences (Fig. 4c and 4d). Exemplary cells labeled 1-3 at long pulse mode and #4-6 at short pulse mode demonstrated differential PI uptake and Ca^2+^ signaling after ultrasound exposure. Cell #1 and #6 both exhibited no membrane poration but mild Ca^2+^ signaling that reached a lower peak value and at a later timing than the other perforated cells (Fig. 4e and Fig.4f). Interestingly, though the final PI uptake in cell #2-3 (long pulse mode) was around 5 times larger than that in cell #4-5 (short pulse mode) (Fig. 4f), the peak values of Ca^2+^ signaling of the latter (1.2-1.5) were larger than those of the former cells (0.45-0.75) (Fig. 4e). This could be attributed to the Ca^2+^ indicator leakage from excess membrane poration for cell #2-3 in the long pulse mode [51]. We further quantified and compared the percentage of cells showing PI uptake and Ca^2+^ signaling in each independent experiment at both pulse sequences. The percentages of both PI positive cells and Ca^2+^ signaling positive cells were significantly higher at long pulse mode than those at short pulse mode (Fig. 4g-h). Again, we observed a significantly higher total detached area at long pulse mode while no discernable cell detachment was observed at short pulse mode (Fig. 4i). As cellular Ca^2+^ signaling reached peak values before apparent PI uptake and cell detachment occurred, we were able to quantify and compare Ca^2+^ peaks, which was measured to be significantly higher at long pulse mode (Fig. 4j). This could be explained by the higher membrane poration rate that allowed Ca^2+^ influx and more cells showing Ca^2+^ signaling at long pulse mode [41]. It is also important to note that excessive Ca^2+^ loading could be toxic to cells.

Cell detachment may cause localized vascular damage or disruption, while non-uniform and excessive membrane poration and Ca²⁺ signaling can lead to uneven transcellular transport and tight junction disruption. In contrast, the absence of detectable cell detachment, along with more uniform and mild membrane poration and a global Ca²⁺ response, would enable safer BBB opening via promoting transcellular transport and a reversible, widespread tight junction opening across a larger vessel area.

### *In vivo* BBB Opening Indicated by Evans Blue Tracing and Tissue Damage Evaluation

Subsequently, the BBB openings by long-pulse and short-pulse ultrasound were performed on mice. Long or short pulse ultrasound was applied to the right hemisphere of the mouse using the ring transducer. Blue staining of Evans blue appeared in the ultrasound-exposed brain tissue regions under the selected acoustic parameters, indicating successful BBB opening, as shown in Fig. 5a. Fluorescence imaging was conducted on these brain tissues, revealing significant fluorescence signals in the ultrasound-exposed areas, which confirmed that Evans blue had crossed the BBB into the brain tissue. The control group, which was only injected with microbubbles and Evans blue without undergoing ultrasound exposure, did not exhibit this effect. Subsequently, we performed sectioning and microscopic examination using a fluorescence microscope, as depicted in Fig. 5b. The results indicated that long pulses induced a more substantial BBB opening than short-pulse ultrasound, as evidenced by a 1.4-fold higher fluorescence intensity. However, the fluorescence signal in the long-pulse ultrasound regions was uneven, while the fluorescence signal in the short-pulse BBB opening areas was more uniform.

**Figure 5.**
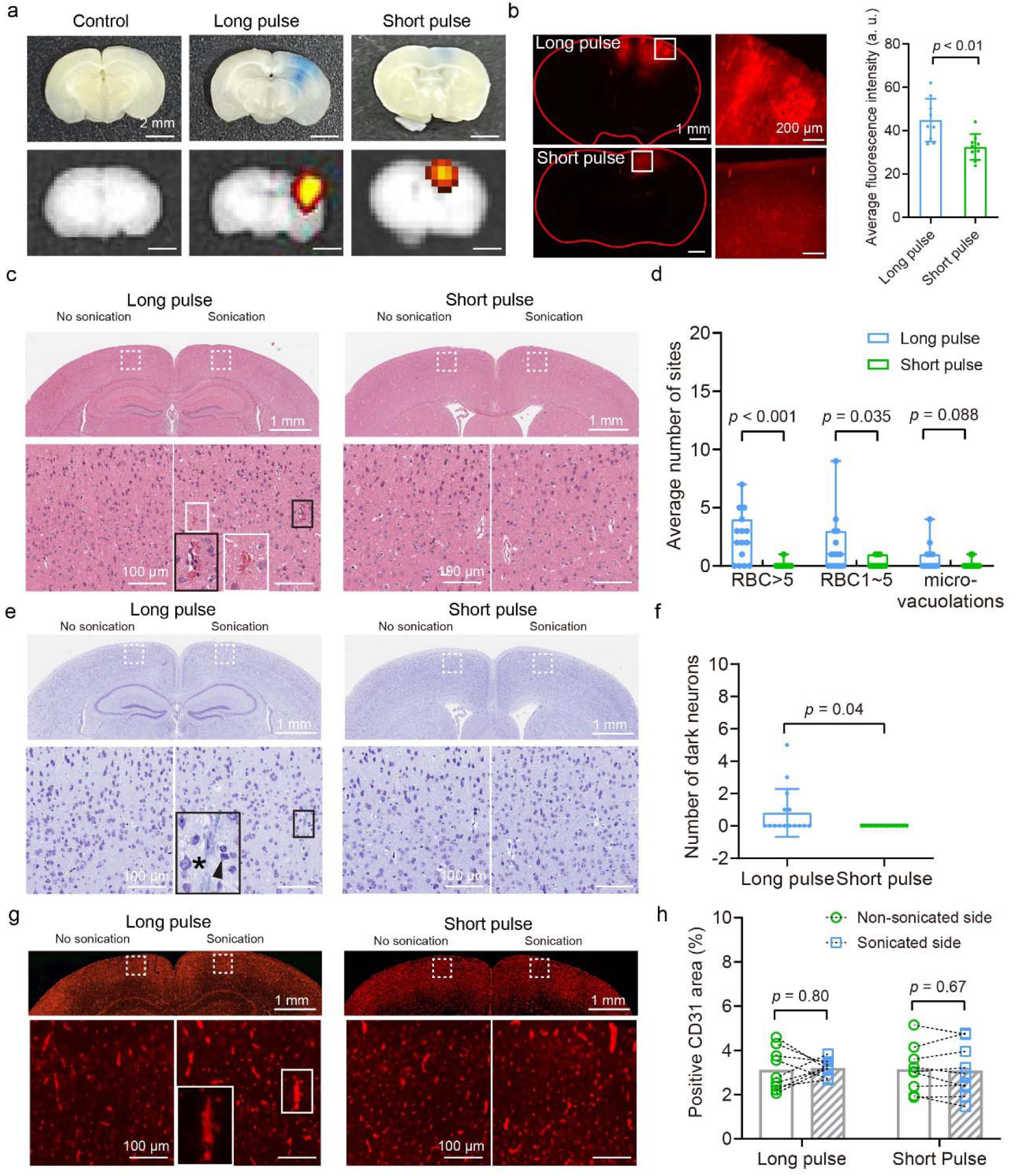
*In vivo* BBB opening assessed through Evans blue tracing and tissue damage evaluation. (a) Evans blue extravasation indicates BBB openings induced by both long-pulse and short-pulse ultrasound. The upper panel displays bright field photographs, while the lower panel shows corresponding fluorescence images. Scale bars: 2 mm. (b) Fluorescence microscopic images illustrating BBB openings resulting from long-pulse and short-pulse ultrasound. The statistical comparison of the average fluorescence intensity is shown on the right. (c) H&E staining images of brain slices exhibiting BBB openings induced by long-pulse and short-pulse ultrasound. (d) Assessment of tissue damage based on H&E staining images. (e) Nissl staining images of brain slices underwent BBB opening due to long-pulse and short-pulse ultrasound. Asterisk: a healthy neuron. Arrowhead: a dark neuron. (f) Quantification of dark neurons surrounding the red blood cell extravasation sites. (g) CD31 immunostaining images of brain slices following BBB opening induced by long-pulse and short-pulse ultrasound. (h) Vessel densities indicated by relative areas of positive CD31 fluorescence in the cortex. H&E, Nissl and CD31 sections are consecutive at the same level. The lower panel images provide enlarged views of the regions outlined by white dashed lines in the upper panel images. Unpaired student t-tests were used for statistical analysis in panel b, d, and f. Paired student t-test was used for statistical analysis in panel h.

H&E, Nissl staining and CD31 immunostaining were performed to assess acute tissue damage. With long pulses at *in situ* 0.5-MPa peak negative pressure, red blood cell extravasation and micro-vacuolations were observed in 2 of 3 brains while only 1 of 3 brains for short-pulse ultrasound (Fig. 5c). The number of sites with more than five extravasated red blood cells induced by long-pulse ultrasound exposure was significantly greater than that caused by short-pulse ultrasound (Fig. 5d). Nissl staining of adjacent brain sections revealed the presence of dark neurons surrounding the red blood cell extravasation site (Fig. 5e-f). However, endothelial immunostaining images showed that positive CD31 signal could be observed at sites with red blood cell extravasation (Fig. 5g). There is no significant difference in vessel density between the contralateral side (no sonication) and sonication side (Fig. 5h).

### *In vivo* Two-photon Imaging of BBB Opening Dynamics by Long-pulse and Short-pulse Ultrasound

The ring transducer designed in this study effectively matches the dimensions of the two-photon water-dipping lens, providing essential experimental conditions for observing the dynamic process of the BBB opening *in vivo*. After calibrating the acoustic field, both long-pulse and short-pulse ultrasound successfully opened the BBB and delivered 500-kDa dextran to the brain tissue. Fig. 6a presents the dynamic two-photon image sequences of the BBB opening using long and short pulse ultrasound, with a total ultrasound exposure duration of 120 seconds (Movie S8 and Movie S9). Compared to long pulses, the short-pulse ultrasound required a longer latency before the BBB began to open. Quantitative analysis revealed that the latency under long-pulse ultrasound was 30.0 ± 16.2 s. In comparison, under short-pulse ultrasound, it was 97.7 ± 34.7 s, with a significant difference (*p* < 0.001) (Fig. 6b). Additionally, from Fig. 6a, we can see that dextran diffused more widely after extravasating from the capillaries under long-pulse conditions. In contrast, the diffusion under short-pulse conditions was more concentrated. As shown in Fig. 6c, long-pulse ultrasound resulted in a larger diffusion area for dextran (11.5 ± 2.5 ×10^3^ µm^2^). In contrast, the diffusion area under short-pulse conditions was only 38.3% of that for long pulses (4.4 ± 1.1 ×10^3^ µm^2^, *p* < 0.001).

**Figure 6.**
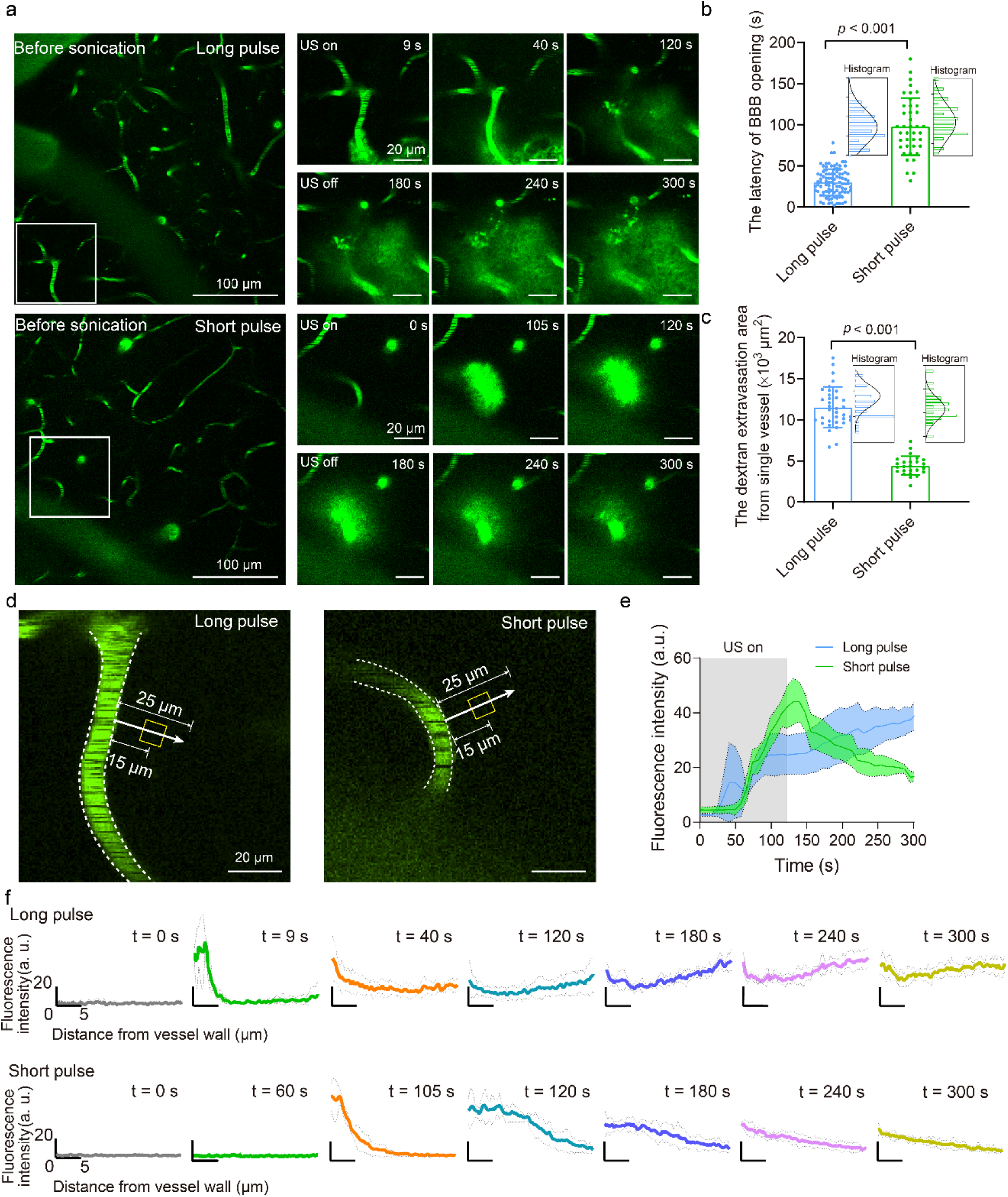
Two-photon time-serial image sequences of the BBB opening using long- and short-pulse ultrasound modes with ultrasound exposure durations of 120 seconds. (a) Time-serial fluorescence images showing the cerebrovasculature before, during, and after sonication at long-pulse and short-pulse modes. The image acquisition lasted 300 seconds. The right-hand side images provide enlarged views of the regions outlined by white boxes in the left-hand side images. (b) The latency of BBB opening induced by long-pulse and short-pulse ultrasound. (c) The calculated average dextran extravasation area from a single vessel at long-pulse and short-pulse modes from each image field. (d-e) Quantification of fluorescence intensity changes with time during dextran diffusion under long-pulse and short-pulse ultrasound. A rectangular region measuring 15 µm perpendicular to the vessel wall was selected. (f) Fluorescence intensity measurements along a path 25 µm from the outer wall of the vessel (indicated by the arrows) under long-pulse and short-pulse ultrasound.

Next, we selected a square region measuring 15 µm perpendicular to the vessel wall, as shown in Fig. 6d, to assess changes in fluorescence intensity over time. As illustrated in Fig. 6e, there was a significant increase in fluorescence intensity within this region during the 120 seconds of sonication, indicating substance diffusion. After sonication, dextran diffusion induced by long-pulse ultrasound continued for an additional 180 seconds throughout the recording, while diffusion from short-pulse ultrasound exhibited a gradual decline. We then measured the fluorescence intensity along a path 25 µm from the outer wall of the vessel. Fig. 6f shows that dextran gradually diffused distally over time under long-pulse ultrasound. However, after 120 seconds, in areas distant from the selected vessel, the fluorescence intensity began to increase, suggesting that the observed dextran leakage was likely due to the opening of the BBB in nearby vessels. In contrast, the fluorescence intensity gradually decreased after a brief diffusion period under short-pulse conditions, with no distal enhancement trend. This indicates that the diffusion characteristics of substances crossing the BBB differ depending on the ultrasound pulse conditions.

### *In vivo* two-photon imaging of endothelial cell loss by long-pulse ultrasound

Due to our observations of red blood cell extravasation in H&E-stained sections following long-pulse ultrasound treatment, we utilized transgenic mice (Tek-iCre: Ai 14) expressing tdTomato fluorescence in endothelial cells for *in vivo* two-photon imaging. Although the number of mice available for this study was limited, we were fortunate to observe a phenomenon where endothelial cell loss occurred during the BBB opening process induced by long-pulse ultrasound (Fig. 7 and Movie S10). The ultrasound emission setup and parameters used were consistent with those described above.

**Figure 7.**
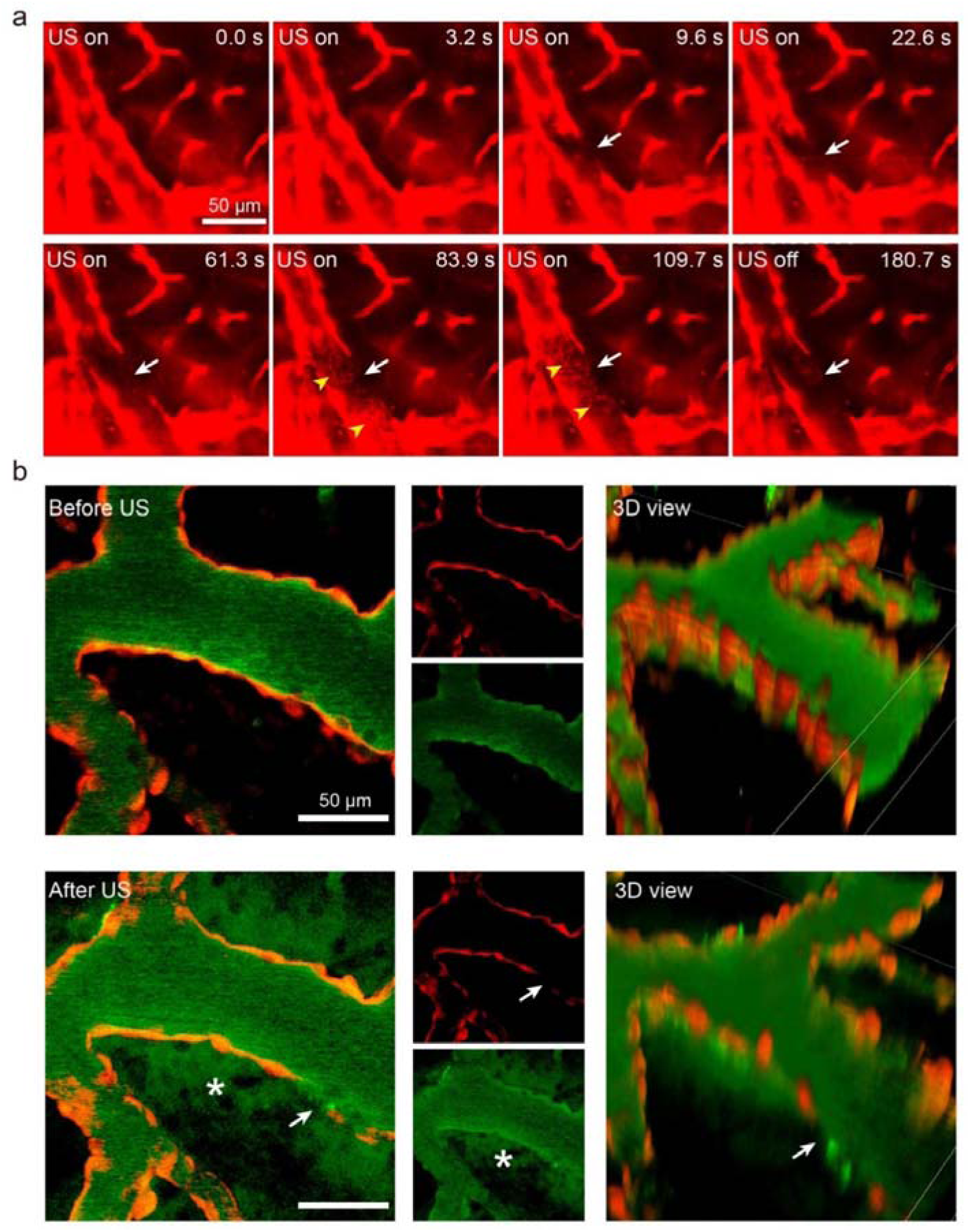
*In vivo* two-photon imaging of endothelial cell loss during long-pulse ultrasound-induced BBB opening in two Tek-iCre: Ai 14 mice. (a) Time-serial imaging reveals a significant loss of tdTomato fluorescence from endothelial cells (white arrows) at 9.6 seconds from ultrasound application, with no recovery detected at 180.7 seconds. Transient flow of sediment-like debris (yellow arrowheads) within the vasculature is observed at 83.9 seconds, subsequently disappearing. (b) Z-stack two-photon fluorescence images of endothelial cells before and after ultrasound treatment demonstrate the loss of fluorescence signal (white arrows) coinciding with the opening of the BBB (white asterisks). Three-dimensional reconstructions enhance visualization of the endothelial cell response at the exact vascular location, highlighting the correlation between long-pulse ultrasound exposure and endothelial cell fluorescence loss. Scale bars: 50 µm.

Figure 7a illustrates that at 9.6 s after ultrasound initiation, there was a noticeable loss of fluorescence signal from the endothelial cells, and this signal did not recover even at 180.7 seconds post-sonication. Notably, at 83.9 s, we observed sediment-like debris flowing through the blood vessels, which disappeared subsequently, implying possible endothelial detachment. Figure 7b presents z-stack two-photon fluorescence scanning images of the endothelial cells before and after the ultrasound-induced BBB opening. By reconstructing three-dimensional images, we gained a clearer vision of, at the same vascular location, the loss of endothelial cell fluorescence signal induced by the long-pulse ultrasound, highlighting the correlation between long-pulse ultrasound BBB opening and endothelial cell loss. In contrast, at the sites of FITC-dextran leakage induced by short-pulse ultrasound, the endothelial cell fluorescence signal remained consistent with that at the initial time point (Fig. S9).

## Discussion

We have developed cross-scale model systems that enable direct observation of bubble dynamics and cellular bioeffects under flow conditions in vessel-mimicking microchannels. These systems also allow us to monitor BBB opening via intravital two-photon imaging in mice and assess tissue damage through histological examination. At an acoustic pressure of 0.5 MPa, long-pulse ultrasound induced stronger cavitation activity, characterized by a more pronounced reduction in bubble number counted and sustained bubble oscillation with cyclic jetting. This was accompanied by topical cell detachment and extensive sonoporation, which correlated with endothelial cell loss and tissue damage. *In vivo* experiments further demonstrated that long-pulse mode enabled significantly higher drug delivery doses and faster BBB opening speeds. In contrast, short-pulse ultrasound produced milder bubble dynamics with a more uniform bubble distribution. This led to reversible and more evenly distributed sonoporation, along with apparent calcium signaling that may facilitate tight junction opening. These findings explain the more even drug delivery and faster BBB recovery observed *in vivo* under short-pulse mode.

Acoustic radiation forces, also known as primary Bjerknes forces, can induce translational movement of oscillating microbubbles. These effects become more pronounced with longer acoustic pulses and higher ultrasonic intensities [37, 59]. Additionally, when microbubbles cavitate, they act as secondary sources of ultrasound waves, exerting forces on neighboring bubbles (secondary Bjerknes forces). This leads to bubble clustering and coalescence, as bubbles attract each other when oscillating in phase. At higher acoustic pressures (0.5 MPa) and in long-pulse mode, we observed significant bubble translation, clustering, and coalescence. These phenomena resulted in a substantially greater reduction in bubble number compared to short-pulse mode. They also explain the more uneven microbubble distribution under 0.5 MPa long-pulse ultrasound exposure.

It is well known that microbubbles oscillate in response to incidental ultrasound waves. In our study, a sinusoidal acoustic driving pulse was applied, and assuming small oscillation amplitudes relative to the equilibrium radius, the bubble’s eigenfrequency in a free field can be expressed as:

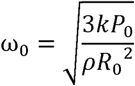

where *k* is the polytropic exponent of the gas-filled microbubble and *P*_0_ is the ambient pressure, *ρ* is the liquid density and *R*_0_ is the equilibrium bubble radius [37]. For heavy gases (*k*=1.1), with *ρ*=1000 kg/m^3^, *P*_0_ = 100 kPa, this yields:

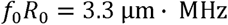

Here *f*_0_ (the Minnaert frequency) serves as a good estimate for coated microbubbles’ resonance frequency [37]. However, bubble oscillation dynamics can be significantly altered near a substrate. In our experimental setup, an air-backing ring ultrasonic transducer transmitted acoustic waves downward into the microchannel, generating acoustic radiation forces along the ultrasound beam direction. These forces pushed microbubbles toward the microchannels’ bottom walls, where cells were located. This enhanced microbubble-cell interaction and facilitated bubble oscillation near the bottom wall. A recent study demonstrated that lipid-coated microbubbles with a perfluorobutane gas core exhibit a resonant radius of ∼3.5 µm near a rigid substrate under 1 MHz acoustic excitation. Notably, cyclic jetting—a phenomenon requiring minimal pressure when the bubble reaches its resonant radius—was observed under these conditions [43]. In our experiments, we similarly detected cyclic jetting during stable oscillations of microbubbles with equilibrium radii of 3.5–4 µm, aligning well with these findings rather than predictions based on the Minnaert frequency. Furthermore, our observations revealed that microbubbles expanded beyond 1 µm during cyclic jetting, consistent with prior report that the first microjet within an ultrasound pulse consistently emerges upon radial expansion of ∼1 µm [43]. This kind of cyclic jetting predominantly occurs in the stable cavitation regime, where bubbles exhibit relatively mild oscillations. Importantly, these microjets significantly enhance cellular permeation by generating stresses at least an order of magnitude greater than previously proposed mechanisms [43]. Initially, most microbubbles in our study had radii well below resonance (<1 µm vs. 3.5 µm). However, after two coalescence events (Fig. 2b, bottom panel, S1 and S2), the bubble approached the resonant radius (∼3.5 µm, with diameter exceeding 7 µm), triggering prolonged stable cavitation and cyclic jetting. In summary, long pulse sequences at 0.5 MPa promoted microbubble translation, clustering, and coalescence. These processes increased the likelihood of resonant oscillation and cyclic jetting, ultimately producing shear stresses an order of magnitude greater than those generated by short pulses.

Recent *in vitro* and *in vivo* studies have demonstrated that transient and reversible tight junction reorganization can mediate safe BBB opening using ultrasound pulses [51, 60]. Our previous work revealed that Ca^2+^ signaling plays a critical role in regulating changes in cell spread area and tight junction remodeling [51, 53]. In the current study, we observed robust yet reversible Ca²⁺ signaling in non-sonoporated or mildly sonoporated cells exposed to short pulses. In contrast, cells subjected to excessive sonoporation during long pulses exhibited weaker Ca²⁺ responses, likely due to leakage of the Ca²⁺ fluorescence indicator. However, both the occurrence rate and average peak amplitude of Ca²⁺ responses remained higher under long-pulse conditions. This enhanced response may be attributed to the significantly increased membrane poration percentage, as membrane pore formation is known to trigger Ca²⁺ influx from the extracellular space [41, 44].

The comparative analysis of long-pulse and short-pulse ultrasound for BBB opening *in vivo* reveals critical insights into their therapeutic potential and mechanistic disparities. Both approaches leverage microbubble-mediated acoustic cavitation to transiently disrupt the BBB, enabling targeted drug delivery [61]. However, their distinct pulse characteristics and energy profiles lead to divergent biological outcomes. Long-pulse ultrasound, characterized by millisecond-scale durations and higher energy deposition, demonstrates superior efficacy in delivering large molecules, such as magnetic resonance imaging (MRI) contrast agents and dextran [62]. For instance, when comparing a rapid short-pulse sequence (13 five-cycle pulses at 1.78 MHz fundamental frequency delivered in 10 ms bursts) with 10-ms long-pulse bursts at 400 kPa peak negative pressure, the latter exhibited an order-of-magnitude higher extravasation of gadobutrol and dextran [62]. Our results further show that long-pulse ultrasound at 500 kPa induced a 1.4-fold greater Evans blue extravasation than short-pulse exposure, consistent with prior reports of enhanced BBB permeability under similar protocols [30]. However, long-pulse ultrasound carries significant safety trade-offs, including elevated risks of erythrocyte extravasation, microhemorrhage, and delayed BBB recovery (>24 hours), which correlate with neuroinflammatory responses such as microglial activation [25, 63]. These adverse effects were thought to stem from inertial cavitation dominance in long pulses, generating broadband emissions and pronounced high-order harmonics / ultra-harmonics, whereas short-pulse sequences exhibit negligible such signatures under identical acoustic conditions [30]. Our *in vitro* observations suggest that prolonged cyclic jetting during stable cavitation may also contribute to these side effects in the absence of inertial cavitation.

Short-pulse ultrasound reduces energy exposure while achieving homogeneous molecular distribution across the BBB, spanning from small to large molecules [30, 31, 48]. Yet, prior studies have not explored the dynamic processes or mechanisms of large-molecule BBB penetration mediated by these modalities at the microscopic level in live brains. Utilizing two-photon intravital imaging, we observed distinct spatiotemporal patterns: short-pulse ultrasound exhibited a threefold longer latency in BBB opening initiation compared to long-pulse protocols. This delay aligns with *in vitro* vessel-mimicking experiments showing slower calcein leakage kinetics under short-pulse conditions, likely due to milder microbubble-endothelium interactions. Notably, cranial window observations revealed endothelial cell detachment and localized fluorescence voids exclusively in long-pulse groups, corroborated by in vitro findings. These microstructural disruptions explain the heterogeneous dextran extravasation and prolonged BBB permeability seen with long-pulse ultrasound.

Time-serial tracking of FITC-dextran diffusion revealed rapid clearance kinetics post short-pulse exposure, contrasting with persistent accumulation under long-pulse conditions. This suggests that long-pulse ultrasound induces irreversible BBB disruption, whereas short-pulse exposure mediates transient, reversible permeabilization. The observed PI uptake and Ca²⁺ signaling in short-pulse cohorts support a reversible sonoporation and tight junction modulation mechanism, as opposed to the irreversible membrane compromise induced by long pulses. These findings elucidate why short-pulse ultrasound facilitates rapid BBB recovery (<10 min), while long-pulse protocols require 24 hours for restoration [30]. The accelerated recovery with short pulses minimizes neurotoxic exposure to blood-derived components and may mitigate inflammatory cascades.

The choice between these modalities hinges on therapeutic objectives. Long-pulse strategies remain essential for delivering bulky agents but require careful parameter optimization to mitigate safety risks. Conversely, short-pulse protocols offer a paradigm shift toward precision BBB modulation, particularly for small-molecule CNS drugs or diagnostic tracers. Future research should explore the molecular weight limits of short-pulse ultrasound by optimizing microbubble dosing or pressure escalation while preserving safety. Additionally, the interplay between pulse repetition frequency and microbubble cavitation dynamics warrants deeper investigation to refine spatiotemporal control over BBB permeability.

The cross-scale model systems developed in this study provide the first comprehensive, real-time microscopic analysis of the dynamics and biophysical mechanisms governing ultrasound-mediated BBB opening with microbubbles. These insights not only deepen our understanding of how to achieve safer and more efficient BBB modulation but also open new avenues for precisely tailored and targeted drug delivery through vascular routes—both to the central nervous system (CNS) and peripheral tissues. By elucidating the key factors controlling BBB permeability, this work lays the foundation for optimized ultrasound protocols with broad therapeutic potential, particularly in treating tumors, neurodegenerative diseases, and other CNS disorders.

## Materials and Methods

### Vessel-Mimicking Microchannel

The vessel-mimicking microchannels were fabricated using the standard soft photolithography technique (please refer to SI for more information). Each microfluidic chip consists of three independent microchannels with dimensions of 200 µm × 100 µm × 17000 µm (width × height × length). Marker patterns were designed to align the x-y plan focus of the ring ultrasound transducer with the microchannels (bottom view in Fig.1a). The height of the Polydimethylsiloxane (PDMS) was kept at 3 mm to allow the ultrasound beam to focus in the microchannels in the z-direction.

### Cell Culture, Preparations and Handling

The bEnd.3 cells (murine brain microvascular endothelial cells.3) were routinely cultured in DMEM medium (Gibco, C11995500BT) supplemented with 10% heat-inactivated fetal bovine serum and 1% penicillin/streptomycin at 37 ℃ with 5% CO_2_. BEnd.3 cells of passage numbers 5-12 were used in this study. We followed previous protocols to seed cells inside the microchannels [41, 64] (please refer to SI for more information). Cells were then cultured with DMEM growth medium for around six hours under a perfusion flow rate of 2 µL/min with a syringe pump (KDS, R462) to form cell monolayer. Before experiments, cells in the microchannels were perfused with DMEM growth medium with 5 µg/ml Hoechst 33342 (Biosharp, BL803A) and 1 μM calcein-AM (Invitrogen, 2525771) or 6 μM Fluo4-AM (Invitrogen, F14201) to label cell nuclei, indicate membrane integrity or calcium signaling, respectively. After incubating at 37°C for 15 minutes in the dark, unloaded dyes were washed with 1X DPBS at 2 µL/min for 1 min. Then the microfluidic chip with cells was mounted on the microscope. The channels were perfused with 1X DPBS (with Ca^2+^) containing microbubbles (diluted 1:20 v/v) and propidium iodide (PI, Thermo Fisher Scientific, P21493) at a target concentration of 100 µg/mL at a flow rate of 75 µL/min using the syringe pump.

### Setup for Bubble Dynamics and Cellular Bioeffects in Microchannels under Ultrasound Exposure with Microbubbles

For the experiments of bubble dynamics, aforementioned home-made polydisperse microbubbles were diluted 1:20 v/v in 1X DPBS and injected into the microchannels at a flow rate of 75 µL/min using the syringe pump on an inverted microscope (Zeiss, Axio observer 7), see Fig.1a. Microbubbles were sonicated with a 1.125 MHz custom-built ring-shaped ultrasound transducer, which was driven by a 50-dB power amplifier (2100 L, Electronics & Innovation, USA). The ring transducer was placed at the top surface of the PDMS chip through high vacuum grease gel (DOW CORNING) and aligned with the microchannels by the aiding alignment markers (bottom panel of Fig. 1a). The ultrasound waveform was generated by a function generator (DG972, RIGOL, China). The acoustic pressure output through the PDMS layer was calibrated with a hydrophone (HNR-0500, Onda Corporation, USA). Two distinct ultrasound sequences were used in this study: long-pulse (One burst per second with 9.09-ms-long pulses for 60 bursts) and short-pulse mode (60 bursts in total with one burst per second, within each burst, the pulses were emitted at a repetition frequency (PRF) of 1 kHz with a pulse length of 100 µs), as shown Fig. 1c. Two different peak rarefactional pressures, 0.25 MPa and 0.50 MPa were used after attenuation through PDMS. Bubble dynamics were recorded through a 63× objective (LD PN 63x/0.75 Corr) with a high-speed camera (Nova S12, Photron) synchronized with the ultrasound burst at 1s intervals with random manual mode (see Fig. S1e for the detailed imaging sequences). The high-speed camera was operated at 25,000 frames per second (fps) and an exposure time of 0.66 μs. For long pulse mode, bubble dynamics for only one burst (9.09ms) could be recorded due to limited memory of the high-speed camera. For the experiments of cellular bioeffects, the high-speed camera was replaced by an sCMOS camera (EDGE 4.2; PCO) to capture the bright field and fluorescent signals of cells through a 5× microscope objective (N-Achroplan 5x/0.15). Concurrent calcein / PI or Fluo-4 / PI imaging with a switching interval ∼1.1 s were performed through automated systems (please refer to Fig. S1f for more information). Cell morphology and cell nuclei before and after ultrasound treatment was characterized with single-shot bright-field images and Hoechst fluorescence imaging, respectively.

### *In vitro* Image Processing and Data Analysis

High speed videos of bubble dynamics were imported into MATLAB (The MathWorks, Natick, MA, USA, academic use) to analyze the bubbles’ size and number. All images were pre-processed with two bilinear interpolations for image upscaling [47]. The centroid and the diameter of individual bubbles in each image frame were then detected by circular Hough transform. The average volume-weighted bubble diameter was calculated for each image frame: 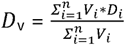 Where *D_i_* is the diameter of the *i* th bubble, *V_i_* is the volume of the *i* th bubble (*V_i_* = 4*π*/3(*D_i_*/2)^3^, and *n* is the total number of bubbles in the image frame. For quantification of the bubble dynamics, relative diameter and number changes of bubbles were defined as 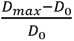 and 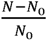, where *D*_0_ is the mean value of *D*_v_ over the 200-μs recording period before ultrasound, and *D*_max_ is the maximum *D*_v_ during the ultrasound exposure, *N*_0_ is the average number of bubbles over the 200 μs recording before ultrasound application, and *N* is the average number of bubbles over the 200 μs imaging time after ultrasound exposure. Briefly, we obtained the total number of cells and their locations with automatic detection of the cell nuclei through the Stardist module of ImageJ 1.54f (NIH, USA) for the Hoechst staining images before ultrasound exposure. Similarly, we detected the number of PI-positive cells before and after ultrasound exposure.

#### Animals

Female C57BL/6J mice were used in this study (6–8 weeks old, 20 ± 2 g; Guangdong Medical Laboratory Animal Center, Foshan, China). All mice were housed in a specific-pathogen-free environment, with sufficient food and drinking water, stable ambient temperature and humidity (23-25 °C, 55 ± 5%), and a 12-h light/dark cycle. Animal care and experiments were approved by the Institutional Animal Care and Use Committee of the School of Medicine at Shenzhen University, China (Protocol #202300169) and followed the ARRIVE guidelines [65]. Three female C57BL/6J mice were randomly assigned to each group to assess BBB opening between short-pulse and long-pulse ultrasound. Eight C57BL/6J female mice underwent short-pulse ultrasound during intravital two-photon imaging experiments, while twelve female mice received long-pulse ultrasound treatment. Additionally, four transgenic mice were generated by crossing Tek-iCre line (Strain No. T003764, GemPharmatech, Nanjing, China) with Ai 14 reporter mice (Strain No. 007914, The Jackson Laboratory, U.S.A., provided by Prof. Woo-ping Ge, Chinese Institute for Brain Research, Beijing). These transgenic mice exhibited strong tdTomato fluorescence in brain endothelial cells, allowing for observation of the effects of long-pulse ultrasound with microbubbles treatment on these cells.

#### Histological Examination and Analysis

Following the sonication procedure, Evans blue, which binds to albumin upon entering the bloodstream, was administered intravenously and circulated for 2 hours. The mice were then profoundly anesthetized and underwent intracardiac perfusion with 4% paraformaldehyde (PFA). Their brains were extracted from the skull and fixed in PFA overnight. Fluorescence images of EB accumulation in the brains were captured using a Xenogen IVIS Spectrum system (IVIS SPECTRUM; PerkinElmer Inc., Waltham, MA, USA) with an excitation wavelength of 430 nm and a 680 nm emission filter. The brains were then embedded in paraffin and sectioned into coronal slices with a thickness of 7 μm. Sections were cut at five levels, each 150 μm apart, with three consecutive sections obtained at each level. These sections underwent hematoxylin and eosin (H&E) staining, Nissl staining, and CD31 immunostaining to evaluate potential microvascular injury and neuronal damage. Endothelial cells were stained using primary rabbit anti-CD31 antibodies (1: 200 overnight, GB113151, Servicebio, Wuhan, China) and secondary Cy3-conjugated goat anti-Rabbit IgG antibodies (1: 300, GB21303, Servicebio, Wuhan, China). H&E and Nissl staining sections were imaged by a digital pathology slide scanner (Aperio CS2; Leica, Germany) using a 20 × objective. Immunofluorescence images were obtained with an inverted fluorescence microscope (Eclipse Ti2, Nikon, Japan) using a 20 × objective. Five H&E-stained sections from each brain were evaluated blindly, focusing on the number of areas with more than five extravasated red blood cells and micro-vacuolations [31]. The presence of dark neurons was assessed in five Nissl-stained sections per brain. Additionally, endothelial cell damage was evaluated through CD31 immunostaining images of three sections.

#### The Blood-brain Barrier Opening *in vivo* via Ultrasound with Microbubbles Observed by Two-photon Intravital Imaging

As shown in Fig. 1d, the ring transducer was engineered to match the geometric specifications of the 40× water-dipping objective of an upright two-photon microscope (A1R MP, Nikon, Japan). The acoustic pressure output of the ring transducer through a cover glass was calibrated using a hydrophone (HNR-0500, Onda Corporation, USA) in an Acoustic Intensity Measurement System (Onda Corporation, USA). The mouse, equipped with a cranial window (Fig. 1d), was secured in a custom-built stereotactic setup that held the titanium holder and positioned on the two-photon microscope stage. The ring transducer was placed within the holder, fixed above the cranial window’s cover glass. The objective was to dip into the degassed water inside the ring transducer. The mouse was anesthetized with 1.5% isoflurane (RWD Life Science, China), with the body temperature maintained through a heating pad set at 37.5 °C. FITC-labeled dextran (Molecular weight: 500 kDa, CAS No. 60842-46-8, Maokang Biotechnology Co., Ltd., Shanghai, China) was administered via a tail vein catheter. Then, the vascular bed was imaged by the two-photon microscope. A region of interest was selected and imaged as the background before sonication. Then, a bolus of microbubbles diluted in 0.1 mL saline was injected into the catheter of the mouse at a dose of 2 × 10^5^/g of body weight, followed by a saline flush. Approximately 15 s after the injection of microbubbles, ultrasound was emitted by the ring transducer for 120 s, with 0.50-MPa peak-rarefactional pressure (measured in water with the attenuation of the glass coverslip). The long-pulse and short-pulse parameters matched those of the *in vitro* experiments. Time-serial scanning of two-photon imaging was performed once the ultrasound emission started and lasted 300 seconds. A 40× water-dipping objective lens (CFI Apochromat NIR 40X W, 0.80 NA, 3.5 mm WD) was used for imaging with 920-nm excitation and 525-nm emission filter. 12-bit images were captured at 1024 × 1024 pixels with a lateral resolution of 0.31 μm/pixel. Through the cranial window chamber, brain vasculature could be imaged down to the depth of 600 μm (Fig. S2).

## Acknowledgments

This research was supported by the start-up funding from Shenzhen Bay Laboratory, the Natural Science Foundation of Guangdong Province (Nos. 2023A1515010649 and 2024A1515012591), the Guangdong Provincial Pearl River Talents Program 2023QN10X235 and the National Natural Science Foundation of China (No. 12474458 and 12204322). The authors would like to thank Jianpeng Wei for assistance in data analysis of the measured acoustic field. They thank Professor Woo-ping Ge from the Chinese Institute for Brain Research in Beijing for generously providing transgenic mice. They also thank the Instrumental Analysis Center of Shenzhen University (Lihu Campus) for their assistance with two-photon intravital imaging.

## Notes

**Competing Interest Statement:** The authors declare no conflict of interest.

### Competing Interest Statement

The authors have declared no competing interest.

## References

1. Daneman, R. and A. Prat, The blood-brain barrier. Cold Spring Harb Perspect Biol, 2015. 7(1): p. a020412.

2. Choi, Y.K. and K.W. Kim, Blood-neural barrier: its diversity and coordinated cell-to-cell communication. BMB Rep, 2008. 41(5): p. 345–52.

3. Wu, D., et al., The blood-brain barrier: structure, regulation, and drug delivery. Signal Transduct Target Ther, 2023. 8(1): p. 217.

4. Andreone, B.J., et al., Blood-Brain Barrier Permeability Is Regulated by Lipid Transport-Dependent Suppression of Caveolae-Mediated Transcytosis. Neuron, 2017. 94(3): p. 581–594.e5.

5. Pardridge, W.M., Drug transport across the blood-brain barrier. J Cereb Blood Flow Metab, 2012. 32(11): p. 1959–72.

6. Cai, Y., et al., Advances in BBB on Chip and Application for Studying Reversible Opening of Blood-Brain Barrier by Sonoporation. Micromachines (Basel), 2022. 14(1).

7. Meng, Y., et al., Safety and efficacy of focused ultrasound induced blood-brain barrier opening, an integrative review of animal and human studies. J Control Release, 2019. 309: p. 25–36.

8. Park, E.J., et al., Ultrasound-mediated blood-brain/blood-tumor barrier disruption improves outcomes with trastuzumab in a breast cancer brain metastasis model. J Control Release, 2012. 163(3): p. 277–84.

9. Ting, C.Y., et al., Concurrent blood-brain barrier opening and local drug delivery using drug-carrying microbubbles and focused ultrasound for brain glioma treatment. Biomaterials, 2012. 33(2): p. 704–12.

10. Jordão, J.F., et al., Antibodies targeted to the brain with image-guided focused ultrasound reduces amyloid-beta plaque load in the TgCRND8 mouse model of Alzheimer’s disease. PLoS One, 2010. 5(5): p. e10549.

11. Kinoshita, M., et al., Targeted delivery of antibodies through the blood-brain barrier by MRI-guided focused ultrasound. Biochem Biophys Res Commun, 2006. 340(4): p. 1085–90.

12. Chen, P.Y., et al., Focused ultrasound-induced blood-brain barrier opening to enhance interleukin-12 delivery for brain tumor immunotherapy: a preclinical feasibility study. J Transl Med, 2015. 13: p. 93.

13. Baseri, B., et al., Activation of signaling pathways following localized delivery of systemically administered neurotrophic factors across the blood-brain barrier using focused ultrasound and microbubbles. Phys Med Biol, 2012. 57(7): p. N65–81.

14. Samiotaki, G., et al., Enhanced delivery and bioactivity of the neurturin neurotrophic factor through focused ultrasound-mediated blood--brain barrier opening in vivo. J Cereb Blood Flow Metab, 2015. 35(4): p. 611–22.

15. Thévenot, E., et al., Targeted delivery of self-complementary adeno-associated virus serotype 9 to the brain, using magnetic resonance imaging-guided focused ultrasound. Hum Gene Ther, 2012. 23(11): p. 1144–55.

16. Weber-Adrian, D., et al., Gene delivery to the spinal cord using MRI-guided focused ultrasound. Gene Ther, 2015. 22(7): p. 568–77.

17. Burgess, A., et al., Targeted delivery of neural stem cells to the brain using MRI-guided focused ultrasound to disrupt the blood-brain barrier. PLoS One, 2011. 6(11): p. e27877.

18. Alkins, R., et al., Focused ultrasound delivers targeted immune cells to metastatic brain tumors. Cancer Res, 2013. 73(6): p. 1892–9.

19. Anastasiadis, P., et al., Localized blood-brain barrier opening in infiltrating gliomas with MRI-guided acoustic emissions-controlled focused ultrasound. Proc Natl Acad Sci U S A, 2021. 118(37).

20. Baek, H., et al., Clinical Intervention Using Focused Ultrasound (FUS) Stimulation of the Brain in Diverse Neurological Disorders. Front Neurol, 2022. 13: p. 880814.

21. Blesa, J., et al., BBB opening with focused ultrasound in nonhuman primates and Parkinson’s disease patients: Targeted AAV vector delivery and PET imaging. Sci Adv, 2023. 9(16): p. eadf4888.

22. Gasca-Salas, C., et al., Blood-brain barrier opening with focused ultrasound in Parkinson’s disease dementia. Nat Commun, 2021. 12(1): p. 779.

23. Abrahao, A., et al., First-in-human trial of blood-brain barrier opening in amyotrophic lateral sclerosis using MR-guided focused ultrasound. Nat Commun, 2019. 10(1): p. 4373.

24. Novell, A., et al., A new safety index based on intrapulse monitoring of ultra-harmonic cavitation during ultrasound-induced blood-brain barrier opening procedures. Sci Rep, 2020. 10(1): p. 10088.

25. Kovacs, Z.I., et al., Disrupting the blood-brain barrier by focused ultrasound induces sterile inflammation. Proc Natl Acad Sci U S A, 2017. 114(1): p. E75–e84.

26. Carpentier, A., et al., Clinical trial of blood-brain barrier disruption by pulsed ultrasound. Sci Transl Med, 2016. 8(343): p. 343re2.

27. Shin, J., et al., Focused ultrasound-mediated noninvasive blood-brain barrier modulation: preclinical examination of efficacy and safety in various sonication parameters. Neurosurg Focus, 2018. 44(2): p. E15.

28. Pouliopoulos, A.N., et al., Safety evaluation of a clinical focused ultrasound system for neuronavigation guided blood-brain barrier opening in non-human primates. Sci Rep, 2021. 11(1): p. 15043.

29. Jordão, J.F., et al., Amyloid-β plaque reduction, endogenous antibody delivery and glial activation by brain-targeted, transcranial focused ultrasound. Exp Neurol, 2013. 248: p. 16–29.

30. Morse, S.V., et al., Rapid Short-pulse Ultrasound Delivers Drugs Uniformly across the Murine Blood-Brain Barrier with Negligible Disruption. Radiology, 2019. 291(2): p. 459–466.

31. Morse, S.V., et al., Liposome delivery to the brain with rapid short-pulses of focused ultrasound and microbubbles. J Control Release, 2022. 341: p. 605–615.

32. Choi, J.J., et al., Noninvasive and localized neuronal delivery using short ultrasonic pulses and microbubbles. Proc Natl Acad Sci U S A, 2011. 108(40): p. 16539–44.

33. Lim Kee Chang, W., et al., Rapid short-pulses of focused ultrasound and microbubbles deliver a range of agent sizes to the brain. Sci Rep, 2023. 13(1): p. 6963.

34. Pouliopoulos, A.N., et al., Rapid short-pulse sequences enhance the spatiotemporal uniformity of acoustically driven microbubble activity during flow conditions. J Acoust Soc Am, 2016. 140(4): p. 2469.

35. Choi, J.J., et al., Noninvasive and localized blood-brain barrier disruption using focused ultrasound can be achieved at short pulse lengths and low pulse repetition frequencies. J Cereb Blood Flow Metab, 2011. 31(2): p. 725–37.

36. López-Aguirre, M., et al., The road ahead to successful BBB opening and drug-delivery with focused ultrasound. J Control Release, 2024. 372: p. 901–913.

37. Roovers, S., et al., The Role of Ultrasound-Driven Microbubble Dynamics in Drug Delivery: From Microbubble Fundamentals to Clinical Translation. Langmuir, 2019. 35(31): p. 10173–10191.

38. Marmottant, P. and S. Hilgenfeldt, Controlled vesicle deformation and lysis by single oscillating bubbles. Nature, 2003. 423(6936): p. 153–6.

39. Ohl, C.-D., et al., Sonoporation from Jetting Cavitation Bubbles. Biophysical Journal, 2006. 91(11): p. 4285–4295.

40. Prentice, P., et al., Membrane disruption by optically controlled microbubble cavitation. Nature Physics, 2005. 1(2): p. 107–110.

41. Li, F., et al., Dynamics and mechanisms of intracellular calcium waves elicited by tandem bubble-induced jetting flow. Proc Natl Acad Sci U S A, 2018. 115(3): p. E353–e362.

42. Sankin, G.N., F. Yuan, and P. Zhong, Pulsating Tandem Microbubble for Localized and Directional Single-Cell Membrane Poration. Physical Review Letters, 2010. 105(7): p. 078101.

43. Cattaneo, M., et al., Cyclic jetting enables microbubble-mediated drug delivery. Nature Physics, 2025.

44. Fan, Z., et al., Spatiotemporally controlled single cell sonoporation. Proc Natl Acad Sci U S A, 2012. 109(41): p. 16486–91.

45. Helfield, B., et al., Biophysical insight into mechanisms of sonoporation. Proc Natl Acad Sci U S A, 2016. 113(36): p. 9983–8.

46. Zhao, X., A. Wright, and D.E. Goertz, An optical and acoustic investigation of microbubble cavitation in small channels under therapeutic ultrasound conditions. Ultrason Sonochem, 2023. 93: p. 106291.

47. Le, D.Q., V. Papadopoulou, and P.A. Dayton, Effect of Acoustic Parameters and Microbubble Concentration on the Likelihood of Encapsulated Microbubble Coalescence. Ultrasound in Medicine & Biology, 2021. 47(10): p. 2980–2989.

48. Pouliopoulos, A.N., et al., Rapid short-pulse sequences enhance the spatiotemporal uniformity of acoustically driven microbubble activity during flow conditions. The Journal of the Acoustical Society of America, 2016. 140(4): p. 2469–2480.

49. Chen, X., et al., Dynamic Behavior of Microbubbles during Long Ultrasound Tone-Burst Excitation: Mechanistic Insights into Ultrasound-Microbubble Mediated Therapeutics Using High-Speed Imaging and Cavitation Detection. Ultrasound Med Biol, 2016. 42(2): p. 528–538.

50. De Cock, I., et al., Ultrasound and microbubble mediated drug delivery: Acoustic pressure as determinant for uptake via membrane pores or endocytosis. Journal of Controlled Release, 2015. 197: p. 20–28.

51. Lin, J., et al., Reversible Ca(2+) signaling and enhanced paracellular transport in endothelial monolayer induced by acoustic bubbles and targeted microbeads. Ultrason Sonochem, 2025. 112: p. 107181.

52. Silvani, G., et al., Reversible Cavitation-Induced Junctional Opening in an Artificial Endothelial Layer. Small, 2019. 15(51): p. e1905375.

53. Li, F., et al., Mechanically induced integrin ligation mediates intracellular calcium signaling with single pulsating cavitation bubbles. Theranostics, 2021. 11(12): p. 6090–6104.

54. Shi, J., et al., Faster calcium recovery and membrane resealing in repeated sonoporation for delivery improvement. Journal of Controlled Release, 2022. 352: p. 385–398.

55. Memari, E., et al., Fluid flow influences ultrasound-assisted endothelial membrane permeabilization and calcium flux. Journal of Controlled Release, 2023. 358: p. 333–344.

56. Shen, Y., et al., Delivery of DNA octahedra enhanced by focused ultrasound with microbubbles for glioma therapy. J Control Release, 2022. 350: p. 158–174.

57. Li, F., et al., Oscillate boiling from microheaters. Physical Review Fluids, 2017. 2(1): p. 014007.

58. Yuan, F., C. Yang, and P. Zhong, Cell membrane deformation and bioeffects produced by tandem bubble-induced jetting flow. Proc Natl Acad Sci U S A, 2015. 112(51): p. E7039–47.

59. Lentacker, I., et al., Understanding ultrasound induced sonoporation: definitions and underlying mechanisms. Adv Drug Deliv Rev, 2014. 72: p. 49–64.

60. Noel, R.L., et al., Safe focused ultrasound-mediated blood-brain barrier opening is driven primarily by transient reorganization of tight junctions. bioRxiv, 2025: p. 2025.01.28.635258.

61. Batts, A.J., et al., Using a novel rapid alternating steering angles pulse sequence to evaluate the impact of theranostic ultrasound-mediated ultra-short pulse length on blood-brain barrier opening volume and closure, cavitation mapping, drug delivery feasibility, and safety. Theranostics, 2023. 13(3): p. 1180–1197.

62. McMahon, D., L. Deng, and K. Hynynen, Comparing rapid short-pulse to tone burst sonication sequences for focused ultrasound and microbubble-mediated blood-brain barrier permeability enhancement. J Control Release, 2021. 329: p. 696–705.

63. Samiotaki, G. and E.E. Konofagou, Dependence of the reversibility of focused-ultrasound-induced blood-brain barrier opening on pressure and pulse length in vivo. IEEE Trans Ultrason Ferroelectr Freq Control, 2013. 60(11): p. 2257–65.

64. Li, F., et al., A Microfluidic System with Surface Patterning for Investigating Cavitation Bubble(s)-Cell Interaction and the Resultant Bioeffects at the Single-cell Level. J Vis Exp, 2017(119).

65. Percie du Sert, N., et al., The ARRIVE guidelines 2.0: Updated guidelines for reporting animal research. BMC Veterinary Research, 2020. 16(1): p. 242.

