## Supplementary Information for "A bottom-up approach to understanding ultrasound mediated blood–brain barrier opening using long and short pulses"

*Fenfang Li; Lin Ma

**Vessel-Mimicking Microchannel Fabrication.** The vessel-mimicking microchannel was designed by AutoCAD software, and the SU-8-based master mold was fabricated using a standard soft photolithography technique. Polydimethylsiloxane (PDMS, Sylgard 184 Silicone Elastomer Kit, Dow Corning, 10:1 mix ratio, cured at 60 °C for 4 hours) microchannel was cast from the master mold and bonded onto #1 cover glass (25 × 50 $mm^2$) immediately after 50 seconds of treatment in a plasma cleaner (Diener, Zepto one).

**Cell Seeding in Microchannels.** For *in vitro* sonoporation experiments, cells with 90% confluency were trypsinized and resuspended into prewarmed (37 ℃) culture medium to a density of $1x10^7$ /mL before being introduced into the microchannels via 1 ml sterilized syringe. The microchannels were pre-wetted with 1X PBS for 5 min, then coated with 50 μg/mL fibronectin (Roche, 10838039001) for 15 min at 37 ℃ and washed with 1x DPBS (Catalog No. 14040133, Gibco, with $Ca^{2+}$) before cell seeding. Once the injected cells reached the desired density in the microchannels (60-70%), the input and output tubing of the microchannels were clamped, and the device was placed in the incubator for 0.5 hours to allow the cells to adhere under static conditions.

**Preparation and Characterization of Microbubbles.** The lipid components included 1,2-distearoyl-sn-glycero-3-phosphocholine (DSPC) and N-(carbonyl-methoxypolyethylene glycol-2000)-1,2-distearoyl-sn-glycero-3-phosphoethanolamine (DSPE-PEG2000) (Lipoid, Ludwigshafen, Germany) at a molar ratio of 9:1. After fabrication, the concentration and size distribution of the MBs were assessed using a Coulter Counter Multisizer IV (Beckman Coulter Inc., USA), following dilution with Isoton II solution. The morphology of the microbubbles was examined under a microscope (BX-53, Olympus Corporation, Japan).

**Concurrent Fluorescence Imaging through Automated System *in Vitro*.** The exposure out signal of the PCO camera was used to trigger a digital delay generator (DG535, Stanford research systems), which gave the command to trigger the ultrasound system after 20 seconds of baseline fluorescence recording. The fluorescence (FL) images (total recording time 380 s, the exposure time of PI and calcein / Fluo-4 channel were both 100 ms) were recorded through μManager (version 2.0; Open resource), which communicated and coordinated the sequence of operations between the Cool LED fluorescence excitation, microscope filter turret and the PCO camera to achieve concurrent calcein / Fluo-4 and PI imaging with a switching interval ~1.1 s, see Fig. 1D.

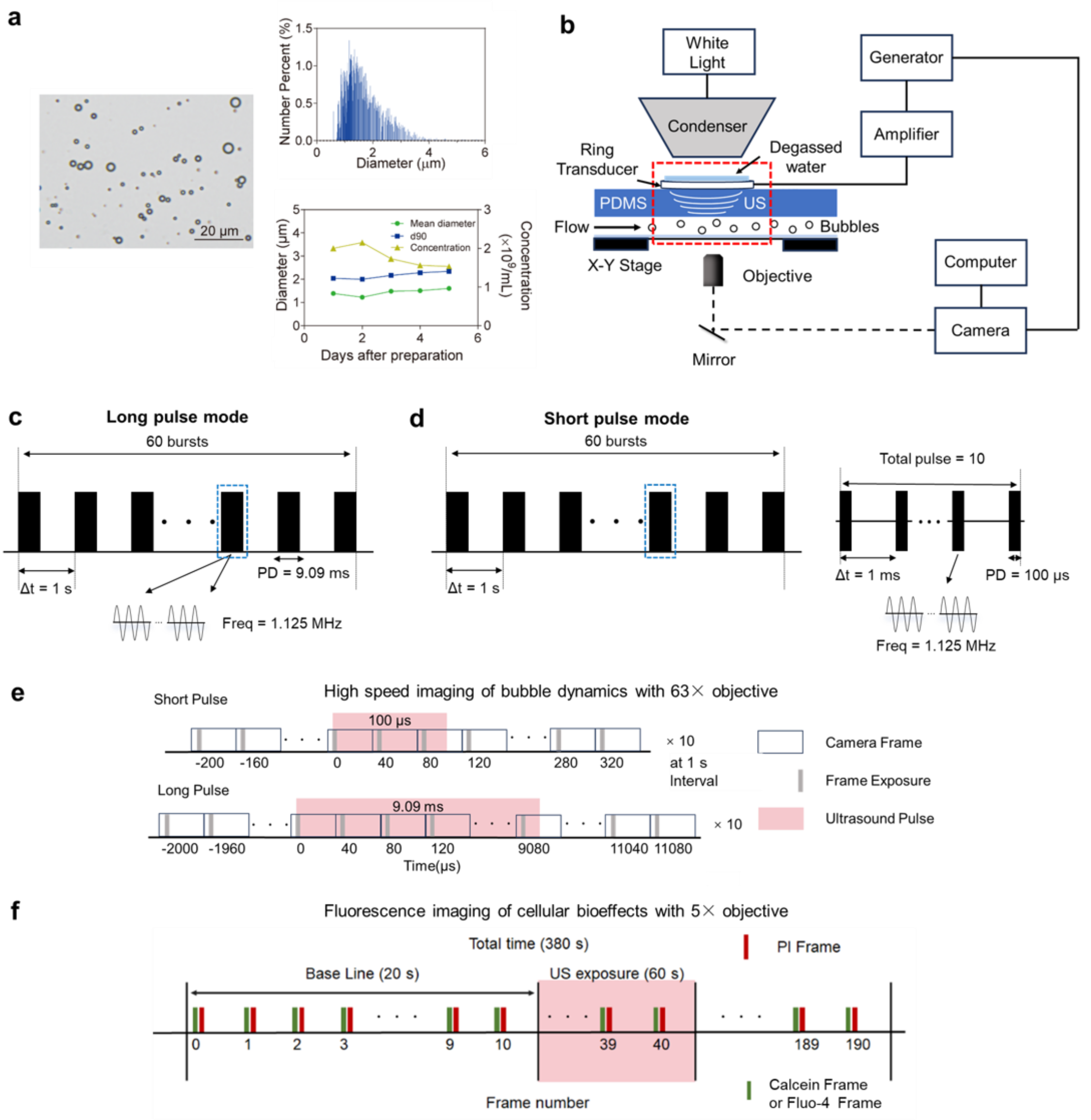


Figure S1. Microbubble characterization, experimental apparatus, acoustic pulse settings and imaging sequences. (a) Characterization of microbubbles: morphology from bright field imaging, size distribution, and stability of microbubbles with a lipid shell and perfluoropropane core, d90: the size below which 90% of the particles fall. (b) Schematic of the detailed experimental setup for ultrasound stimulation system, imaging of bubble dynamics and cellular bioeffects inside microchannels on an inverted microscope through a high-speed camera and an sCMOS camera, respectively. Schematic of the long pulse ultrasound waveforms (c) and short pulse ultrasound sequences (d) used in the experiments. (e) Recording sequences for high-speed imaging of the bubble dynamics and synchronization with ultrasound exposure in short and long pulse mode. (f) Concurrent fluorescent imaging of membrane poration (PI), membrane integrity (calcein) or calcium imaging (Fluo-4), and synchronization with 60 s ultrasound exposure in short and long pulse mode with a 20 s baseline recording.

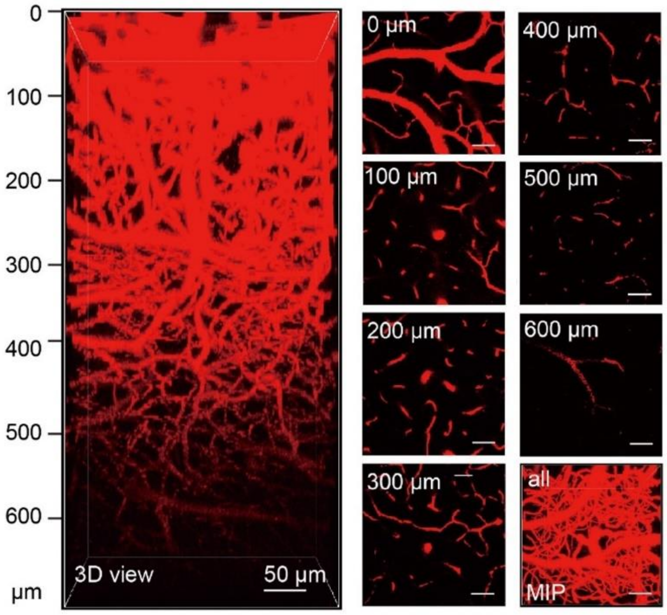


Figure S2. 3D reconstructed images (left) of brain vasculature at different depths from 0 µm to 600 µm (right). MIP: maximum intensity projection of z-stack images. Scale bar: 50 µm.

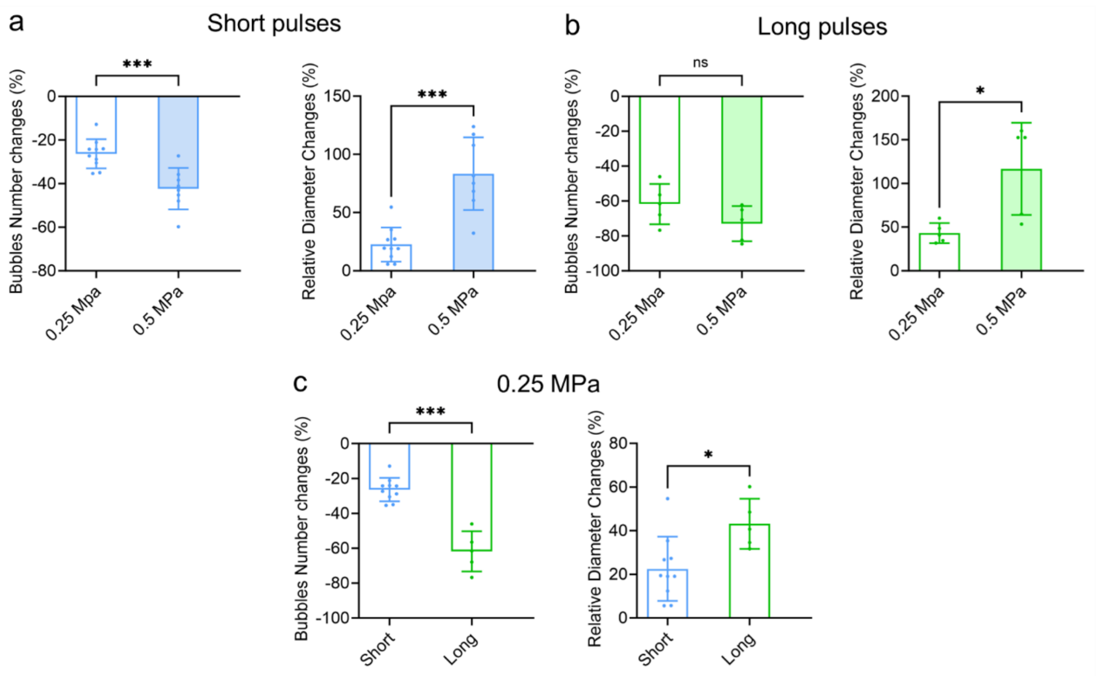


Figure S3. Statistical analysis of bubble numbers and diameter changes under different acoustic parameters. (a) and (b) Comparisons of bubble number reduction and relative bubble size increase at different acoustic pressure in short-pulse and long-pulse mode, respectively. (c) Comparisons of bubble number reduction and relative bubble size increase between short-pulse and long-pulse mode at 0.25 MPa acoustic pressure. Each data point in (a-c) depicts the measurement from an independent experiment. N=8-10 in panel a, N=5 and 5（each group）in panel b, N=5 to 10 in panel c. The student t-test was used for statistical analysis.

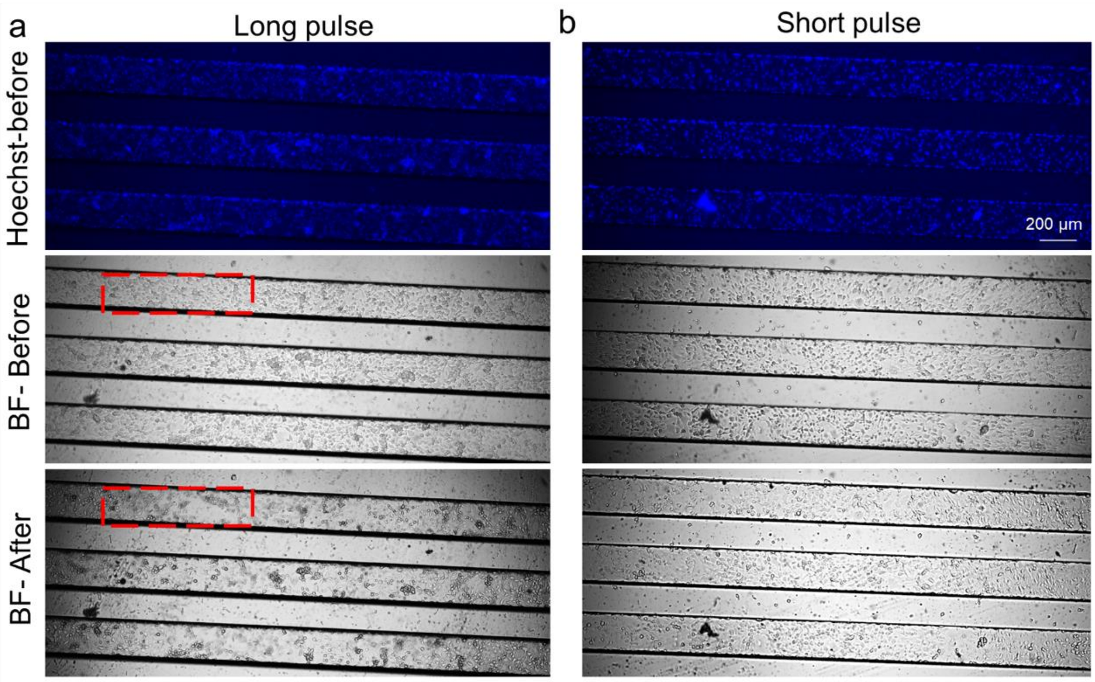


Figure S4. Typical examples showing cell nuclei labeling by Hoechst for cell counting and cell morphology change before and 5 min after 0.5 MPa ultrasound exposure in long-pulse (a) and short-pulse (b) mode. The region in the red box shows cell detachment after long-pulse ultrasound treatment.

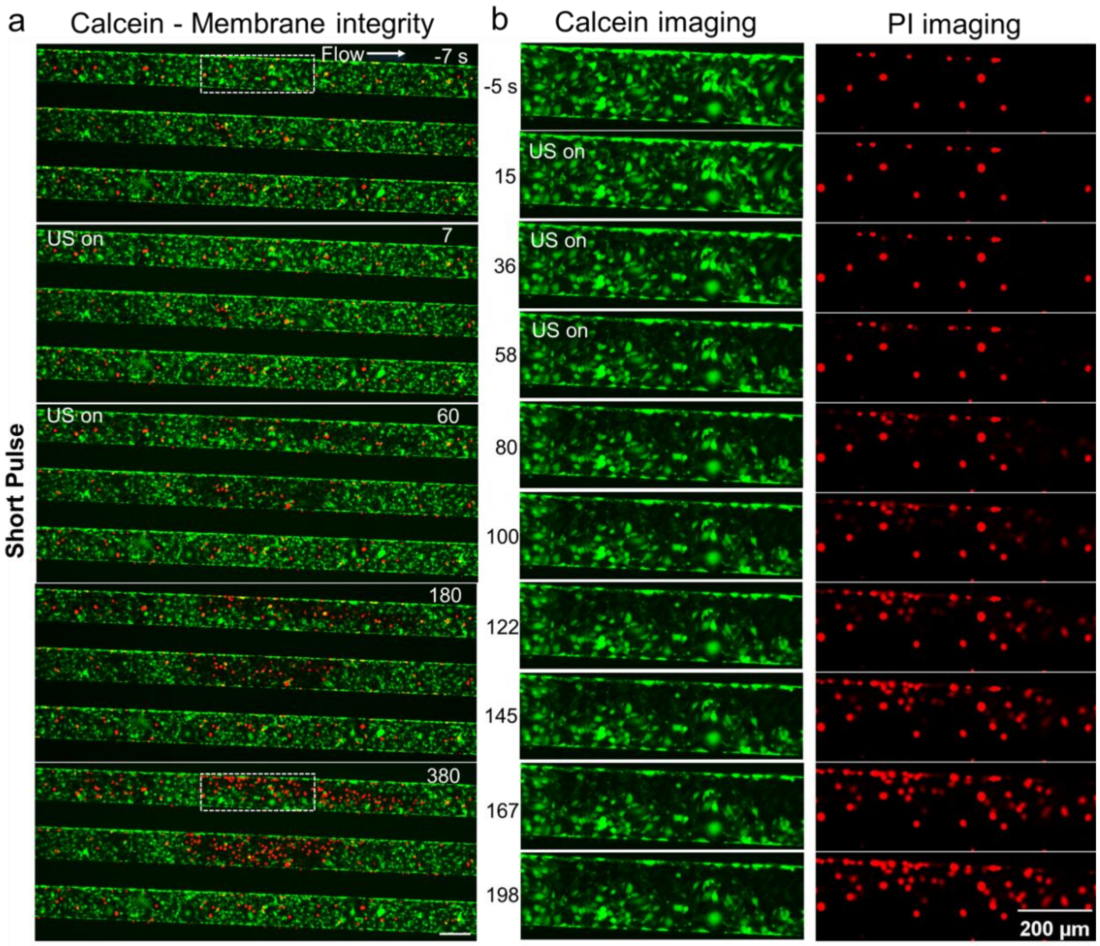


Figure S5. Fluorescence recording of cellular bioeffects at short-pulse mode. (a) Merged image sequence showing the calcein leakage (green) and PI uptake (red) in bEnd.3 cell monolayer in the microchannels before, during (0-60 s), and after 0.5 MPa ultrasound exposure at short-pulse mode. (b) The enlarged view of the dashed box region shows the decrease of cell membrane integrity in panel a with the split channels of calcein (left) and PI (right) fluorescence image sequences. The two channels share the same time labels and scale bar. All the scale bars denote 200 μm.

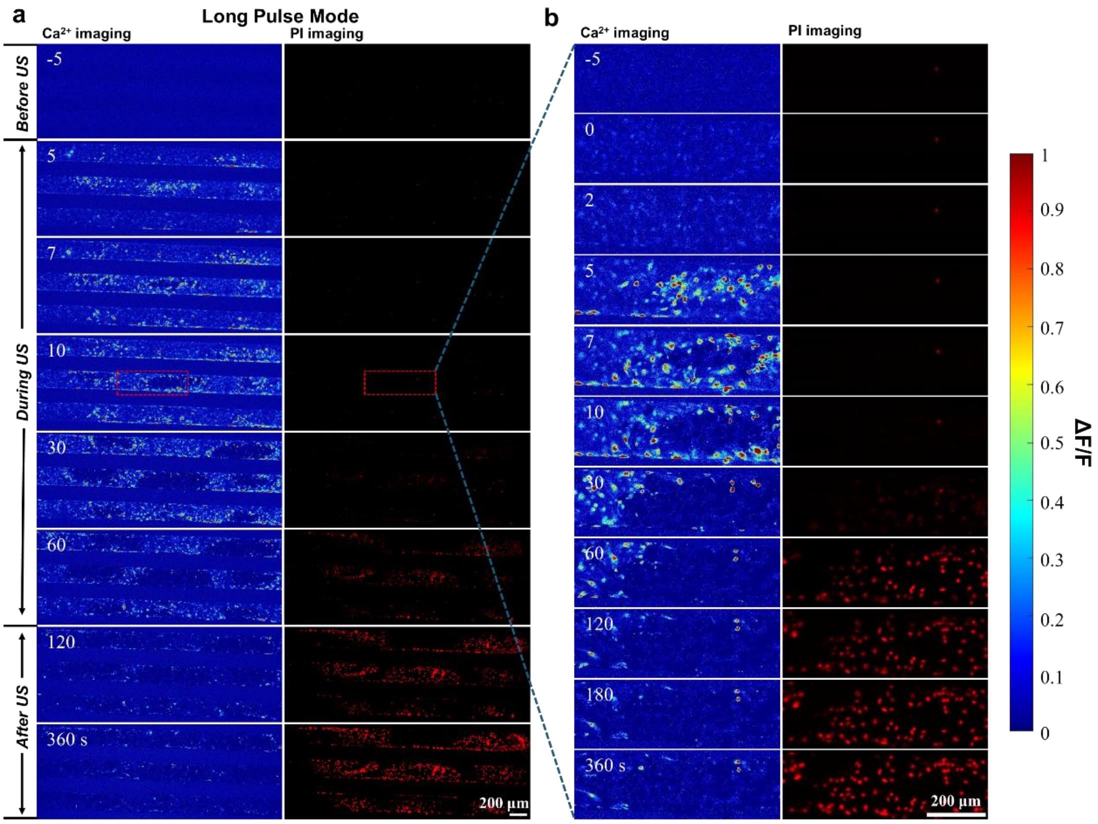


Fig. S6 Characteristics of cellular Ca2+ signaling and membrane poration of bEnd.3 monolayer culture in the vessel-mimicking channel under 0.5 MPa ultrasound exposure at long-pulse mode with further examination of the non-detached region. (a) Merged image sequence showing Ca2+ signaling (Fluo-4, pseudo color) and PI uptake (red) before, during (0-60 s), and after 0.5 MPa ultrasound exposure at long-pulse mode. (b) Enlarged view of the dashed box of a non-detached region in (a) showing the time evolution of Ca2+ signaling (left) and PI uptake (right). The two split channels share the same time labels and scale bar.

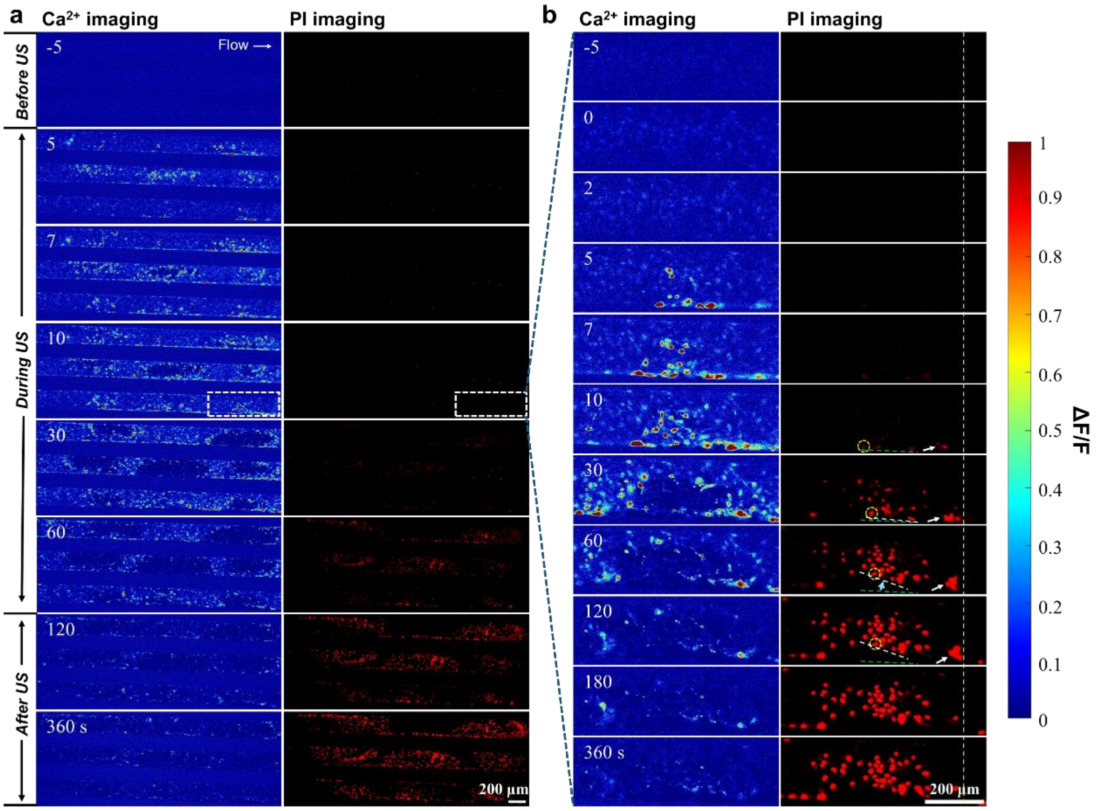


Fig. S7 Characteristics of cellular Ca2+ signaling and membrane poration of bEnd.3 monolayer culture in the vessel-mimicking channel under 0.5 MPa ultrasound exposure at long-pulse mode with further examination of a detached region. (a) Merged image sequence showing Ca2+ signaling (Fluo-4, pseudo color) and PI uptake (red) before, during (0-60 s), and after 0.5 MPa ultrasound exposure at long-pulse mode. (b) Enlarged view of the dashed box of a detached region in (a) showing the time evolution of Ca2+ signaling (left) and PI uptake (right). The dashed lines, the yellow circle and the arrow depict the dynamic displacement of the cells toward the upper right direction. The two split channels share the same time labels and scale bar.

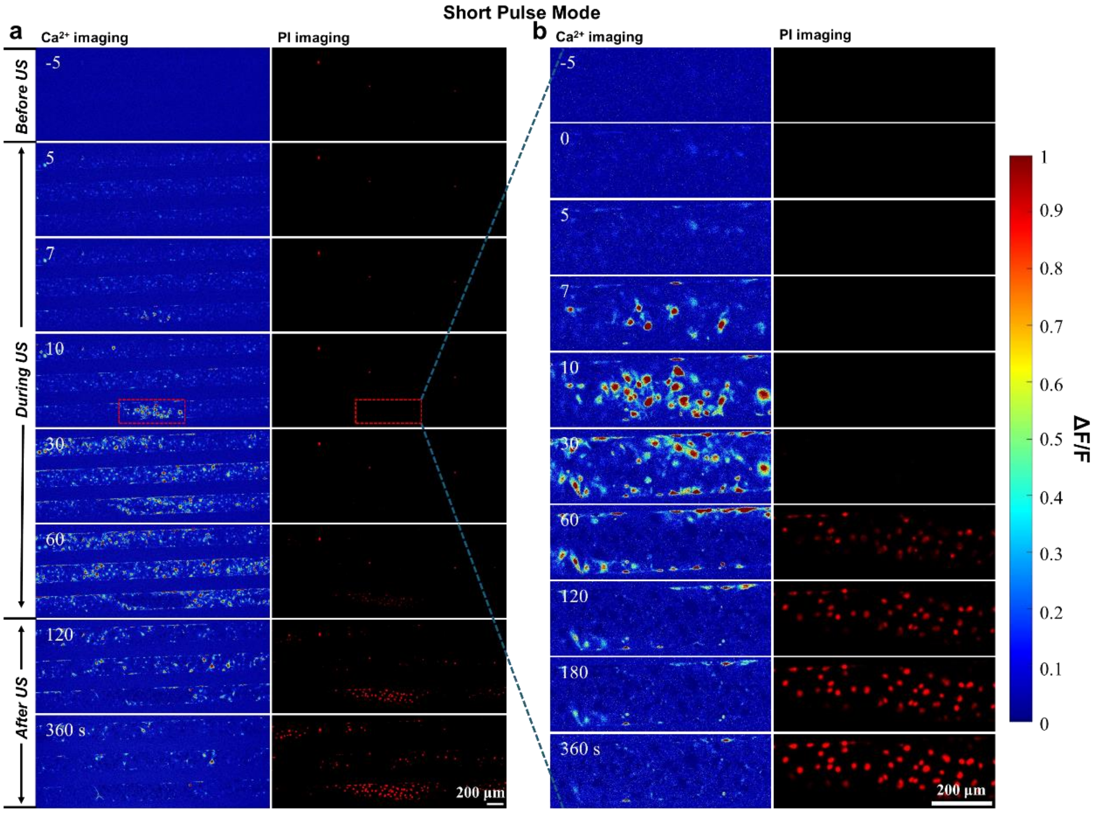


Figure S8. Characteristics of cellular Ca2+ signaling and membrane poration of bEnd.3 monolayer culture in the vessel-mimicking channel under 0.5 MPa ultrasound exposure at short-pulse mode. (a) Merged image sequence showing Ca2+ signaling (Fluo-4, pseudo color) and PI uptake (red) before, during (0-60 s), and after ultrasound exposure. (b) Enlarged view of the dashed box in (a) showing the time evolution of Ca2+ signaling (left) and PI uptake (right), which reveals apparent Ca2+ signaling and evenly distributed sonoporation. The two split channels share the same time labels and scale bar.

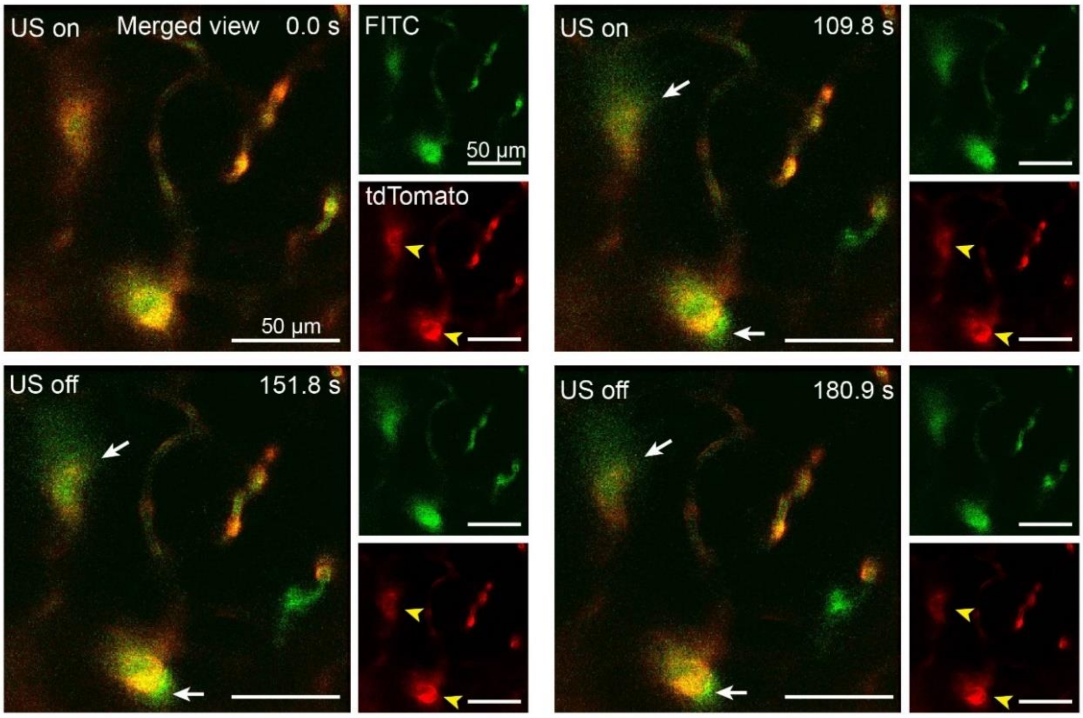


Figure S9. In vivo two-photon imaging during short-pulse ultrasound-induced BBB opening in one Tek-iCre: Ai 14 mice. Time-serial imaging reveals the BBB opening (white arrows) at 109.8 seconds, with no loss of endothelial cells (yellow arrowheads) throughout the recordings.

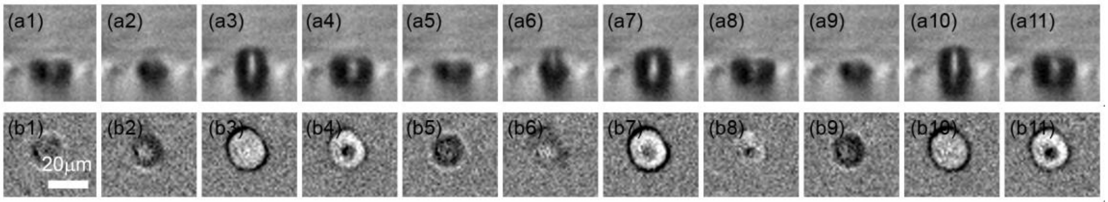


Figure. S10: High speed imaging of stable bubble oscillation with periodic jetting toward the glass substrate from side view (a1-a11) and bottom view (b1-b11). Images have an interframe time of 3:33 ms and an exposure time of 1ms. The formation of a jet propagating towards the substrate during the collapsing phase of the bubble is indicated by the dark spot at the bubble center (frames b4 and b11). Adopted from our previous work[57] (Li, F., et al., Oscillate boiling from microheaters. Physical Review Fluids, 2017. 2(1): p. 014007).

Movie S1. Representative bubble dynamics recorded under short pulse ultrasound of 0.25 MPa, where ultrasound is on between 0-100 μs. Displacement of microbubbles by acoustic radiation force and mild bubble coalescence occur due to secondary Bjerkness force.

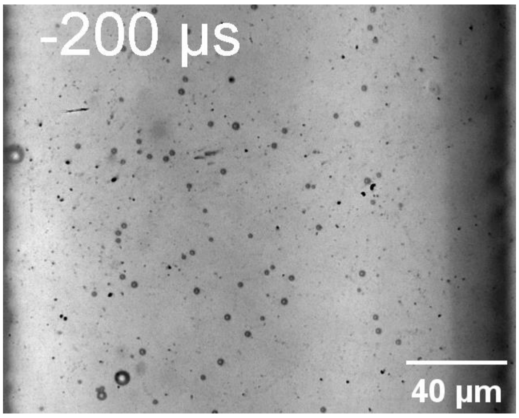


Movie S2. Representative bubble dynamics recorded under short pulse ultrasound of 0.50 MPa, where ultrasound is on between 0-100 μs. Displacement and coalescence of microbubbles become more evident, resulting in larger maximum bubble size and reduced number of bubbles.

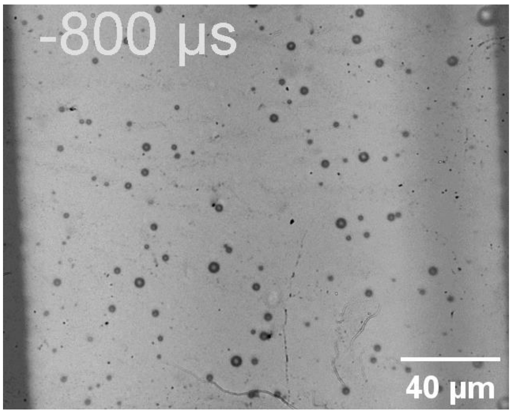


Movie S3. Representative bubble dynamics recorded under long pulse ultrasound at 0.25 MPa acoustic pressure, where ultrasound is on between 0-9090 μs. Displacement and clustering of microbubbles are observed.

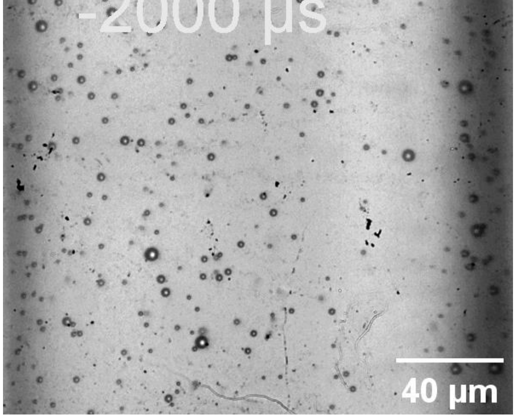

Movie S4. Representative bubble dynamics recorded under long pulse ultrasound at 0.50 MPa acoustic pressure, where ultrasound is on between 0-9090 μs. After bubble displacement, clustering and coalescence in the early stage (0-1000 μs), both stable and inertial cavitation effects occurred in the later stage with bubble oscillation and collapse (1400-9160 μs).

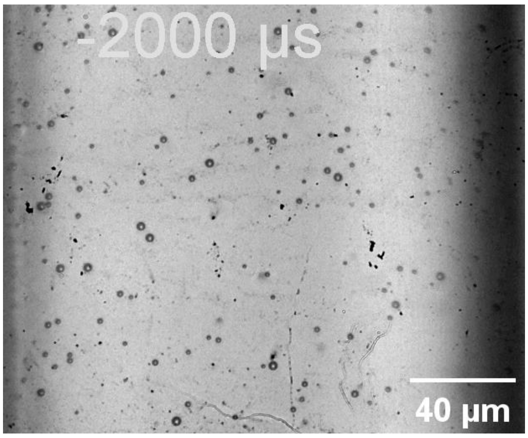


Movie S5. Enlarged view of the stable oscillation and cyclic jetting of the microbubble inside the blue box in Fig. 2b. The bubble kept oscillating from 1240 μs when generated from the coalescence of two bubbles till 4400 μs before it merged with another bubble.

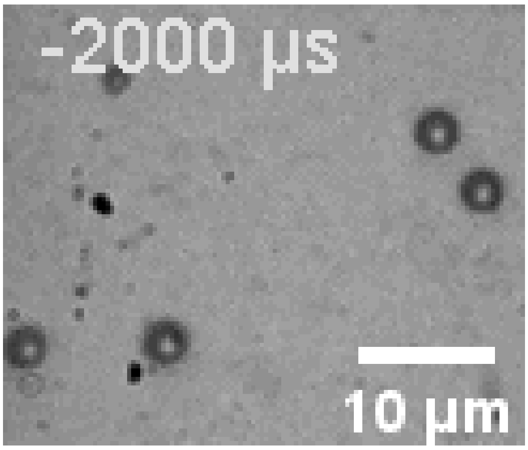

Movie S6. A representative fluorescence recording of cell monolayer response to long pulse ultrasound at 0.5 MPa acoustic pressure. All the viable cells are loaded with calcein (green) before experiment and the green fluorescence decreases with calcein leakage from membrane poration. Cells with membrane poration showed PI uptake (red). Ultrasound is on from 20-80 s here.

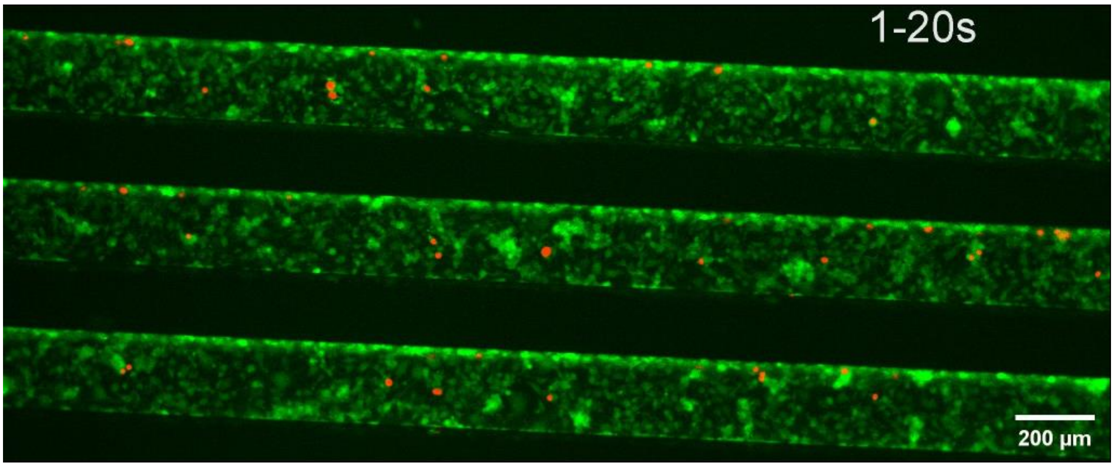


Movie S7. A representative fluorescence recording of cell monolayer response to short pulse ultrasound at 0.5 MPa acoustic pressure. All the viable cells are loaded with calcein (green) before experiment and the green fluorescence decreases with calcein leakage from membrane poration. Cells with membrane poration showed PI uptake (red). Ultrasound is on from 20-80 s here.

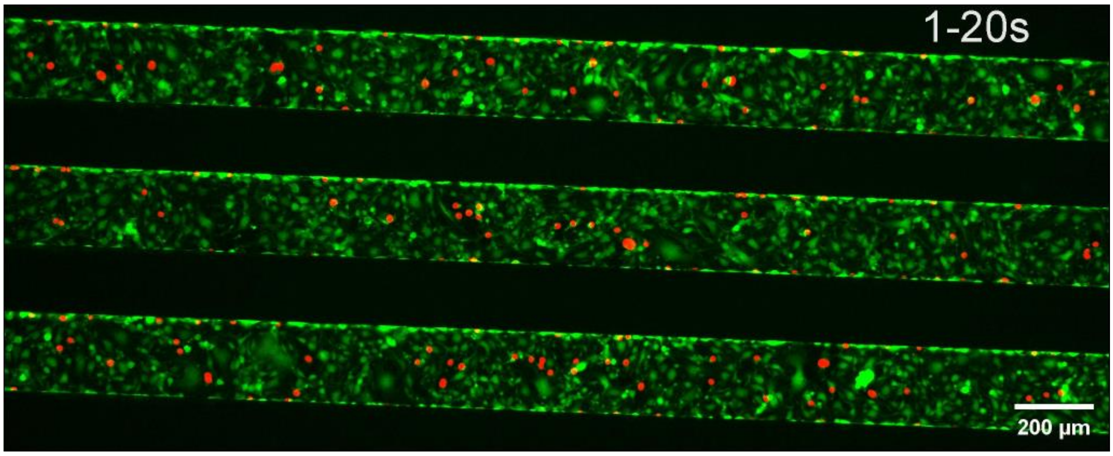

Movie S8. The dynamic two-photon image recording of the BBB opening using long pulse ultrasound, with a total ultrasound exposure duration of 120 seconds.

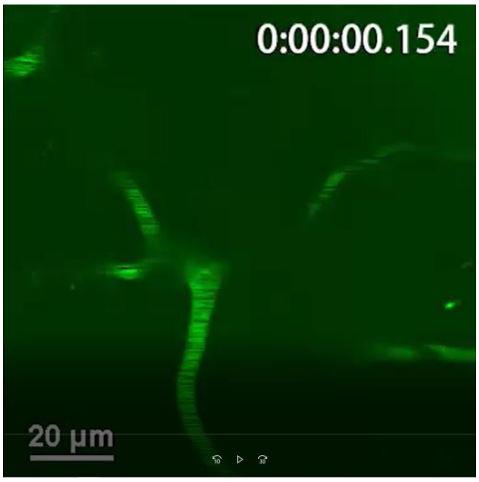


Movie S9. The dynamic two-photon image recording of the BBB opening using short pulse ultrasound, with a total ultrasound exposure duration of 120 seconds.

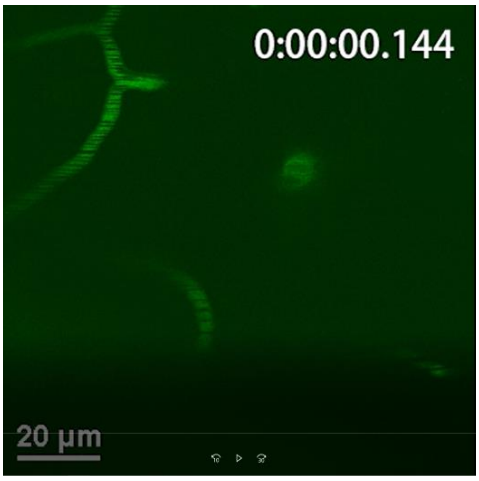


Movie S10. *In vivo* two-photon imaging showing endothelial cell loss during the BBB opening process induced by long-pulse ultrasound at 0.5 MPa acoustic pressure. Transgenic mice (Tek-iCre: Ai 14) expressing tdTomato fluorescence in endothelial cells are used here.

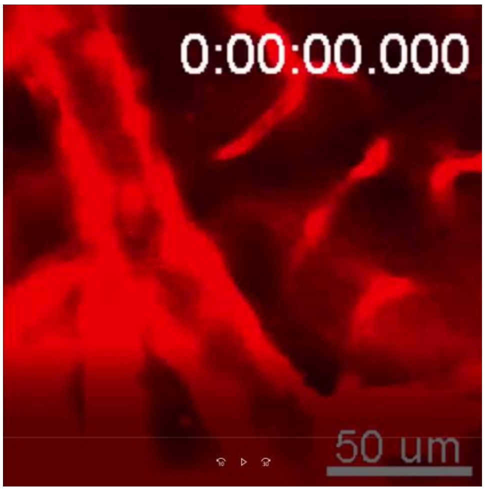
